# Activity and retinoic acid drive hair cell spatial patterning in the zebrafish utricle

**DOI:** 10.1101/2025.11.17.687290

**Authors:** Selina Baeza-Loya, Jo Trang Bùi, David W Raible

## Abstract

The zebrafish vestibular otolith organs, like those of other vertebrate species, are organized into central (striolar) and peripheral (extrastriolar) zones that drive different vestibular circuits. How and when these spatial hair cell patterns develop in the zebrafish is unknown. We determined that early-developing hair cells (<36 hours) expressed both striolar and extrastriolar transcriptomic markers. After 36 hours, these hair cells become specified as extrastriolar hair cells. Later-developing hair cells (>36 hours) mostly develop directly as striolar or extrastriolar. We observed complementary patterns of RA degrading and synthesizing enzymes that colocalize with striolar and extrastriolar hair cells, respectively, indicating evolutionarily conserved molecular signaling. RA treatment during development increased the proportion of extrastriolar and intermediate-type hair cells, indicating that increased RA reduces striolar development. However, in fish with mechanotransduction dysfunction from a *cadherin23* mutation, normal RA patterning is insufficient to finalize the fate of early-born hair cells, which remain transcriptomically unresolved. RA treatment further exacerbates this abnormal patterning. We conclude that hair cell fate, and thus normal zonal patterning, depends on both hair cell activity and the RA gradient.

**Summary statement:** The development of hair cell zonal patterning in the vestibular sensory epithelia depends on a balance of mechanotransduction-driven activity and retinoic acid.

## Introduction

The vestibular organs of the zebrafish inner ear, like those of amniote species, sense linear translations, angular acceleration, and head position in space (reviewed in Baeza-Loya and Raible, 2023). The primary sensory receptors of the inner ear, known as hair cells, detect and transmit motion information through deflection of their eponymous hair bundles. The vestibular organs and associated hair cells are highly conserved across species. These include the three semicircular canals and associated cristae ampullaris (cristae) that sense angular rotation of the head, and the vestibule with associated otolith organs –the utricular and saccular maculae – that detect linear acceleration and gravity.

The organization of the vestibular organs into spatial zones is necessary for a complete vestibular sensory experience and behavioral repertoire. Within the maculae and cristae, hair cells are spatially organized into central and peripheral regions, also referred to as striolar and extrastriolar zones. Hair cells in these central and peripheral zones show conserved functional characteristics across species: hair cells in central striolar zones primarily respond to high frequency stimuli while peripheral extrastriolar hair cells respond to low frequency stimuli (Goldberg, 1991; Tanimoto et al., 2022; Sun et al., 2024). We previously showed that in the zebrafish inner ear, striolar and extrastriolar hair cells had distinct molecular profiles as detected by single cell RNA sequencing, and that these distinct signatures are conserved with mammals (Shi et al., 2023; Smith et al., 2023).

In mouse, retinoic acid (RA) has been shown to drive the development of zones in the otolith epithelia (Ono et al., 2020a, 2020b). RA reduction through the degradation enzyme Cytochrome P450 26b1 (Cyp26b1) is key for the formation of striola (Ono et al., 2020a). RA is generated from its precursor retinal through class 1A3 aldehyde dehydrogenase enzymes (Aldh1a3), which is upregulated in the extrastriolar zone (Ono et al., 2020a, 2020b). Whether RA plays a similar role in the organization of the fish otolith organs is unknown.

In the auditory system, electrical activity originating from hair cells promotes development of the hair cell, the innervating neurons, and the auditory system as a whole (reviewed in Wang and Bergles, 2015). Mechanotransduction-driven activity is also implicated in the maturation of the bioelectrical properties, morphological features, and the synaptic machinery of the hair cell (Corns et al., 2018, Krey et al., 2020, Lee et al., 2021, McQuate et al., 2023). What role mechanotransduction plays in the patterning of hair cells in the vestibular inner ear into zones is poorly understood.

In this study, we assessed the timeline of early zonal development in the larval zebrafish utricle. We trace the cell fate of earliest-developing hair cells to track changes in utricular organization during maturation in the first ten days of development. We characterize the expression pattern of RA degrading and synthesizing enzymes in the utricular epithelia, and tested the effects of RA treatment on hair cell fate. We also assess zonal patterning in fish with dysfunctional mechanotransduction due to a mutation in cadherin23 (*cdh23*). Taken together our results provide evidence for both RA signaling and functional activity in driving the maturation of spatial patterning in the zebrafish inner ear.

## Methods

### Fish maintenance

Experiments were conducted on larval zebrafish between 24 hours post fertilization (hpf) and 5 days post fertilization (dpf), and on juveniles at 10 and 21 dpf. Larvae were raised in E3 embryo medium (14.97 mM NaCl, 500 nM KCl, 42 mM Na_2_HPO_4_, 150 mM KH_2_PO_4_, 1 mM CaCl_2_ dihydrate, 1 mM MgSO_4_, and 0.714 mM NaHCO_3_ at pH 7.2) at 28.5°C in 100 mm petri dishes. Fish intended for juvenile-age experiments were placed on the nursery system at 5 dpf until collection timepoints. Zebrafish experiments and husbandry followed standard protocols in accordance with the University of Washington Institutional Animal Care and Use Committee guidelines.

### Transgenic and mutant lines

The following zebrafish lines were used:

1. *Tg(myo6b:GFP) ^w186^* (Hailey et al., 2017),
2. *Tg(myo6b:nls-Eos) ^w191Tg^* (Cruz et al., 2015) crossed into a nac/roy background (mitfa^w2^; mpv17^a9^; Lister et al., 1999; Ren et al., 2002; White et al., 2008),
3. *cdh23^tj264^* (“*sputnik*”) (Söllner et al. 2004),
4. *cacna1da^tc323d^* (“*gemini*”) (Sidi et al., 2004),
5. *atoh1a^w271Tg^* (Hewitt et al., 2024).

### Photoconversion

Fish larvae with *Tg(myo6b:nls-Eos)* were exposed to UV for ∼8 minutes by being placed in a freezer box lined with aluminum foil (Beaulieu et al., 2024). An iLumen 8 UV flashlight (Amazon) was fixed to the freezer box lid and positioned over the dish.

### Pharmacology

Dechorionated embryos were treated in 24-well cell culture plates starting at 24 hpf with various concentrations of either retinoic acid (RA) (Sigma Aldrich, Cat# R2625), all trans-Retinal (Sigma Aldrich, Cat #R2500), or an equivalent maximum volume of DMSO as a control (≥ 0.1%).

### Fixation

Larvae were fixed in 4% paraformaldehyde (PFA) at 4°C for 12-18 hours. Larvae were then washed three times for 5 minutes each in PBS containing 0.1% Tween20.

### HCR FISH

Hybridization chain reaction (HCR) fluorescent in situ hybridization (FISH) (Molecular Instruments, HCR v3.0) was performed as directed for whole-mount zebrafish embryos and larvae (Choi et al., 2016, 2018, 2020). Briefly, larvae were fixed in 4% PFA at 4°C overnight.

Larvae were washed with PBS, and either dehydrated in methanol and stored at -20°C until use or used immediately. Larvae stored in methanol were rehydrated using a gradation of methanol. Larvae were washed in PBS containing 0.1% Tween20, treated with proteinase K for 25 minutes and postfixed with 4% PFA for 20 minutes at room temperature. For the detection phase, larvae were pre-hybridized with a probe hybridization buffer for 30 minutes at 37°C and then incubated with probes overnight at 37°C. Larvae were then washed with 5x SSC containing 0.1% Tween20 to remove excess probe. For the amplification stage, larvae were preincubated with an amplification buffer for 30 minutes at room temperature and then incubated with hairpins overnight in the dark at room temperature. Excess hairpins were removed by 5x SSCT washes. Hybridized larvae were kept in storage buffer in the dark at 4°C until imaging preparation.

### Imaging preparation and imaging

Fixed fish were mounted on the underside of a coverslip with Fluoromount-G (ThermoFisher, Cat #00-4958-2) mounting media. Specimens were placed on their ventral side in the center of the coverslip with a drop of Fluoromount. A swipe of nail polish on the edge of the coverslip was used to adhere a second coverslip, which was placed on top and gently pushed down to create a seal of mounting media around the specimens. The coverslip sandwiches were left to sit at least 45 minutes at room temperature in the dark while the Fluoromount cured.

Images were captured using a Zeiss LSM-980 with Airyscan 2.0. Z-stacks of inner ear organs were taken using a 40x water objective at intervals of 0.21 µm. All Airyscan processing was performed at standard strength using Zen Blue software (Zeiss, www.zeiss.com). Imaging processing and data analysis were carried out using Fiji (Schindelin et al., 2015).

### Statistical analysis

Statistical analysis and graphing were performed in GraphPad Prism version 10.4.0 (www.graphpad.com) or with Python 3.13. Box plots show mean ± standard deviation (SD). We used an alpha level of 0.05 for all statistical tests. To compare >2 groups, we used a one-way or two-way ANOVA followed by the Bonferroni test for multiple comparisons. We used a chi-square test to assess differences in proportions in hair cell types between wildtype and mutant fish. We used a binomial test to assess probabilities of spatial locations of hair cells on a polar plot. Two-proportion z-tests were used to determine differences in proportion of hair cells between control and drug treated utricles. For significant results, we calculated effect size with bias-corrected Hedges’ g for n<15, Cohen’s d for n≥15 [small effect = 0.2, medium = 0.5, large = 0.8 (Durlak, 2009)], and Cohen’s w for goodness-of-fit tests that used chi-square distributions [small effect = 0.1 – 0.3, medium = 0.3 – 0.05, large = >0.5 (Cohen, 2013)].

### Differential gene expression analysis

Single cell RNA sequencing (scRNA-seq) data of the zebrafish inner ear was mined from Daniocell, a web-based resource of gene expression data from whole-animal wild-type zebrafish embryos and larvae (Farrell et al., 2018; Sur et al., 2023). We utilized the Python-based scVI package to pre-process and create a generative model of count data, followed by visualization, clustering, and differential expression testing with Scanpy package (Wolf et al., 2018; Virshup et al., 2023). Hair cells were selected by expression of *myo6b* and supporting cells by *stm* (Fig S1A-C).

### Computational pipeline for cell classification and spatial analysis

To analyze the spatial organization of utricles across fish, we used a computational approach to classify hair cell types and normalize spatial relationships between hair cells within utricles. 2D orthogonal projection images of 5 dpf utricles were processed as follows: Cells or nuclei were first segmented and labeled as distinct ROIs using Cellpose-SAM (Pachitariu et al., 2025) for cell membranes or StarDist (Schmidt et al., 2018) for cell nuclei, and centroids calculated using regionprops in the scikit-image package 0.25.2 (van der Walt et al., 2014). For each FISH marker, regional intensity was calculated and cells categorized as positive or negative based on thresholding of intensity mean or sum across pixels using regionprops. In Eos experiments, photoconverted cells were manually labeled. For spatial analysis, each image underwent the following transformational pipeline: Cartesian coordinates of the utricle center were calculated and made the new origin (0, 0) to which cells were recentered. The concave hull (Moreira and Santos, 2007) of the recentered utricle was fitted to an ellipse and after converting to polar coordinates, points were rotated so that the major and minor axes of the ellipse became the coordinate axes. Coordinates were then normalized to a unit circle, calculated using the convex hull as the boundary. Two dimensional kernel density estimation was calculated using gaussian_kde from the scipy package version 1.6.2 (Virtanen et al., 2020).

To compare the spatial distributions of hair cells in different conditions, we performed spatial autocorrelation adapted from geographic analysis (Rey et al., 2023). We used the Python Spatial Analysis Library package libpysal v4.13.0 (Rey and Anselin, 2007) to perform join counts and nearest neighbor analysis. KNN weights were calculated with k=7, the lowest value that resulted in no unattached points across all comparisons. Significance was calculated by bootstrap analysis of 9999 permutations.

## Results

### Robust regional patterning of the zebrafish utricle

Striolar and extrastriolar cells are organized into stereotyped zones in the otolith organs by 5 days postfertilization (dpf) (Fig 1) (Smith et al., 2020, 2023; Tanimoto et al, 2022; Shi et al., 2023; Liu et al., 2022; Sun et al., 2024; Beaulieu et al., 2024). We previously identified marker genes for striolar and extrastriolar hair cells using single cell RNA sequencing and hybridization chain reaction fluorescent *in situ* hybridization (HCR FISH) (Shi et al., 2023; Beaulieu et al., 2024) (Fig S1A-E). Expression of *calcium binding protein 1b* (*cabp1b)* is found in hair cells in the medial half of the utricle and the peripheral edges of the saccule, whereas *cabp2b* expression labels hair cells in the lateral half of the utricle and the central strip of the saccule (Fig 1D-E), corresponding to the respective extrastriolar and striolar zones of each epithelium.

**Figure 1:**
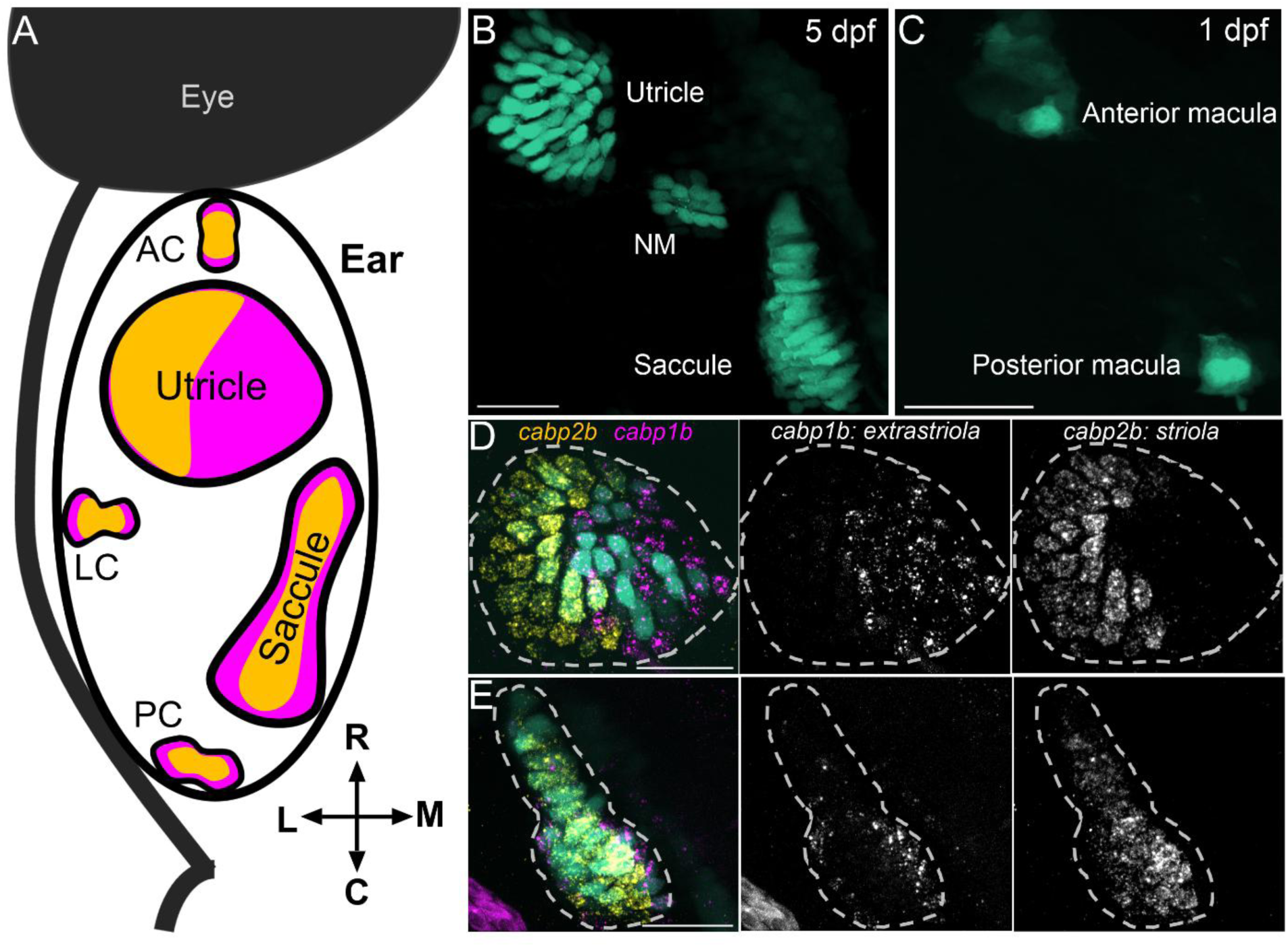
The vestibular epithelia of the zebrafish inner ear have central (striolar) and peripheral (extrastriolar) zones. A) Schematic of a dorsal view of the 5 day postfertilization (dpf) zebrafish inner ear showing five vestibular organs. Yellow areas in the epithelia indicate striolar zones, magenta areas indicate extrastriola. AC: anterior crista, LC: lateral crista, PC: posterior crista. B) Hair cells in the utricle and saccule at 5 dpf in a *Tg(myo6b:GFP)* fish. NM: neuromast. All scale bars = 20 µm. C) Early hair cells in the anterior macula (early utricle) and posterior macula (saccule) at 1 dpf. D & E) At 5 dpf, extrastriolar hair cells are labeled by *cabp1b* (magenta) while striolar hair cells are labeled by *cabp2b* (yellow) in a utricle (D) and saccule (E).

To measure relative spatial organization, we processed 5 dpf utricles by segmenting individual hair cells and classifying them by their molecular identities (Fig 2A). We then projected normalized coordinates of each utricle onto a common framework, retaining the spatial relationships of hair cells within utricles (Fig 2B-C, 10 utricles from 6 fish). We find that regional patterning is highly stereotyped across organs at 5 dpf. At these stages, the vast majority of hair cells express either *cabp1b+* or *cabp2b+*. Out of 703 hair cells, 49% were *cabp1b*+, 47% were *cabp2b*+ (Fig 2B’ – B’’’). Using a binomial test, we assessed whether the proportion of *cabp1b+* or *cabp2b+* hair cells in the medial or lateral half differed from random chance (50%). 302/346 (87%) *capb1b*+ hair cells were in the medial half of the utricle (p < 0.0001), while 297/332 (89%) *cabp2b*+ hair cells were in the lateral half (p < 0.0001).

**Figure 2:**
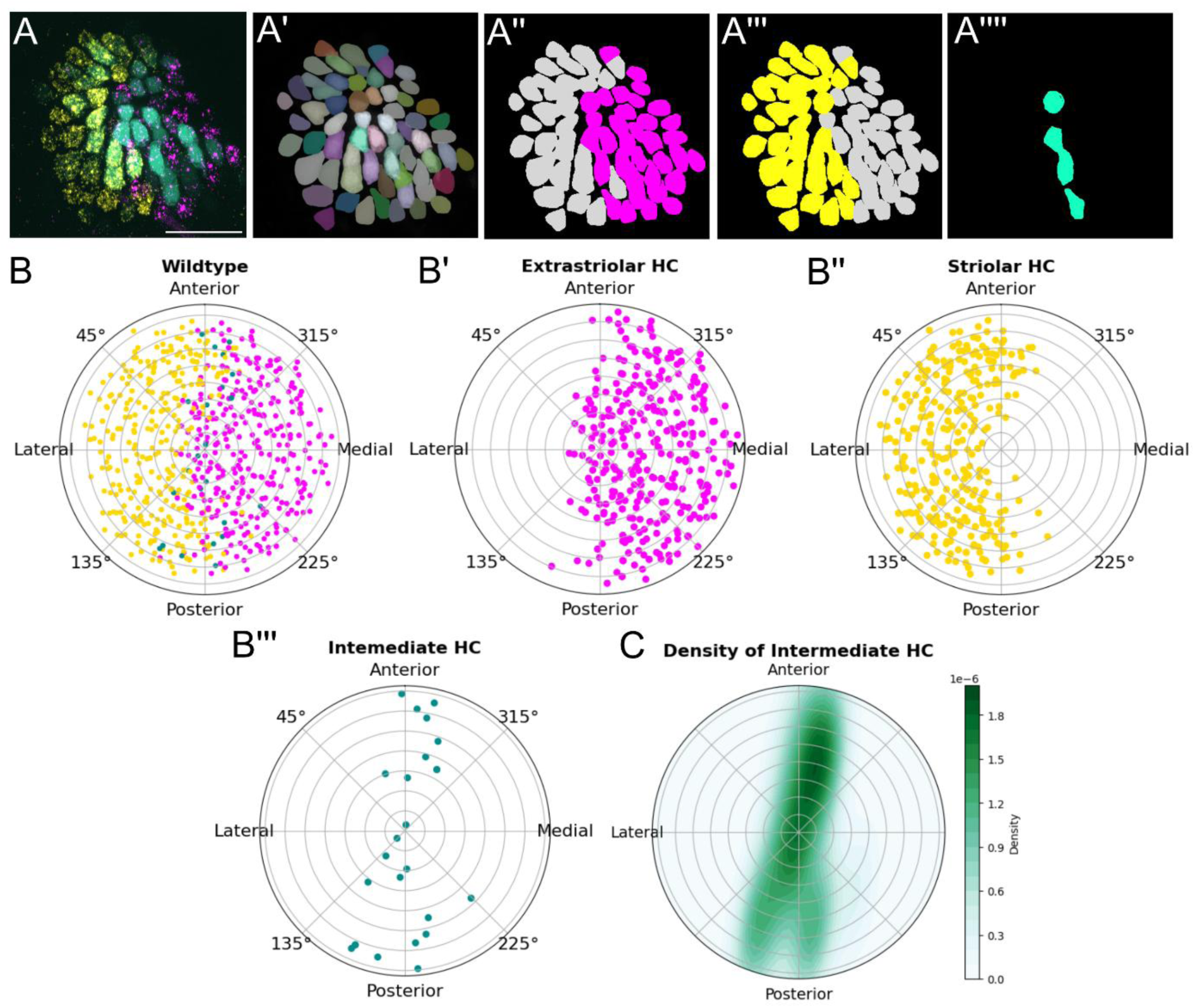
Computational pipeline for cell classification and spatial analysis of utricles. A) Maximum intensity projection images of utricles were processed with Cellpose-SAM to label individual cells with ROI masks (A’). The original max projection images and the cell mask file were processed with an image analysis script to classify cells as either *cabp1b* (A’’), *cabp2b* (A’’’), or both (A’’’’). B) Summary data from wildtype utricles at 5 dpf (n = 10 ears, 6 fish). *Cabp1b*+ hair cells were reliably in the extrastriolar (medial) half (B’), while *cabp2b*+ hair cells were in the striolar (lateral) half (B’’). Intermediate hair cells were located along the zonal boundary (B’’’). C) Density plot of intermediate hair cells from B’’’ shows organization along the A-P pole, coinciding with zonal boundary.

We observed a small fraction (4%) of cells that were positive for both *cabp1b* and *capb2b* probes, indicating they have trancriptomic markers for both hair cell subtypes (Fig 2A, D-E). We call this double-labeled subtype an “intermediate” hair cell. We find that these intermediate hair cells lie along the striolar/extrastriolar boundary (Fig 2B-C). We tested whether the proportion of hair cells in the quarter section of the plot along the boundary (defined as 0 to 22.5°, 202.5° to 337.5°, 337.5° to 360°) differed from random chance (25%). 18 out of 25 cells (72%) of cells were in this range and the proportion was significantly different than chance (binomial test, p < 0.0001).

### Otolith organ striolar and extrastriolar zones are established early in development

We next sought to determine the developmental time course of regional patterning in the otolith organs. Hair cells in the utricle and saccule develop around 24 hours postfertilization (hpf) (Haddon and Lewis, 1996; Tanimoto et al., 2011) (Fig 1C). To determine spatial identity, we again turned to *cabp1b* and *capb2b* expression. Evidence from morphological and functional studies suggests striolar hair cells are more mature relative to extrastriolar hair cells (Liu et al., 2022; Tanimoto et al., 2022). Thus, we hypothesized that early-developing hair cells would have striolar characteristics. To our surprise, utricular and saccular hair cells that develop before 30 hpf were of the intermediate type, positive for both *cabp1b* and *capb2b* probes (Fig 3A, D-E). When we assessed hair cell identity at 36 hpf, we found all hair cells in the utricle were still intermediate. However, by 48 hpf, the utricle primarily had *capb1b*+ hair cells whereas the saccule had mostly *capb2b*+ hair cells (Fig 3D-E). At 5 dpf, the relative proportions of *cabp1b*+ (extrastriolar) to *capb2b*+ (striolar) was close to equal in the utricle; this pattern persisted at 10 dpf (Fig 3D-E).

**Figure 3:**
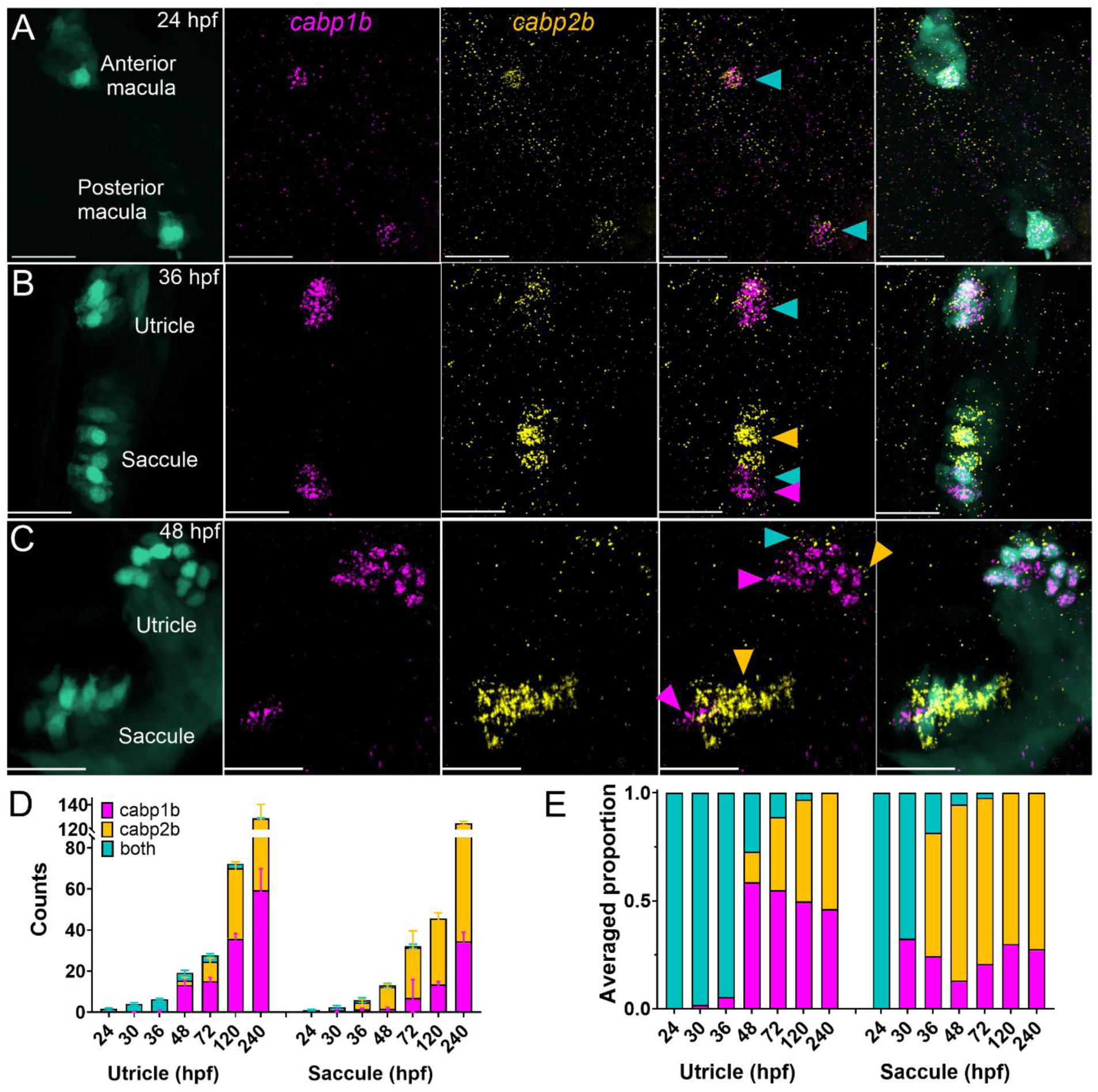
Identification of otolith organ hair cell types during larval development. A – C) Dorsal views of *cabp1b* and *cabp2b* HCR FISH in *Tg(myo6b:GFP)* at (A) 24, (B) 36, and (C) 48 hpf. Hair cells developed before 48 hpf are labeled by both probes (blue arrowheads). These specify into either striolar (yellow arrowheads) or extrastriolar (magenta arrowheads) hair cells during development. Scale bars = 20 µm. D) Hair cells are added in the first 10 days in both the utricle and saccule in an asynchronous and asymmetric manner. Utricle: n = 8 (24 hpf); 12 (30), 8 (36), 13 (48), 9 (72), 13 (120), 12 (240); Saccule: 8 (24), 9 (30), 7 (36), 13 (48), 9 (72), 3 (120), 2 (240). E) Relative proportions of hair cell subtypes change during larval development.

Similar patterns of development are observed in the cristae (Figure S2). The hair cells appeared around 50 hpf in these organs, all of which were *cap2b*+ (Fig S2A). At 54 hpf, some intermediate-type hair cells were detected in all three cristae, and by 72 hpf, *capb1b*+ cells were also observed (Fig S2B-C). By 5 dpf (Fig S2B), hair cells were either *cabp1b*+ or *cabp2b*+ in the three cristae, as previously reported (Beaulieu et al., 2024).

### Hair cell identity changes over the course of early development in the utricle

The reduction in the number of intermediate cells over development suggests that their fates resolve with time. To test this idea, we tracked the identity change of early-developing hair cells using *Tg(myo6b:nls-Eos)* transgenic zebrafish (Beaulieu et al., 2024), which express the photoconvertible protein Eos in hair cell nuclei. Eos exhibits a permanent green to red photoconversion during UV light exposure. In this experiment, we exposed fish to UV light at 36 hpf, and then fixed them at 2, 5, or 10 dpf, and imaged the utricles after HCR FISH. Later-developing hair cells (after 36 hpf) only have green Eos whereas early-developing hair cells (before 36 hpf) retain red converted protein. At 2 dpf, most photoconverted hair cells are still intermediate (*cabp1b+cabp2b+*) (Fig 4A). Later-developing hair cells (not photoconverted) were either intermediate or striolar (*cabp2b*+). At 5 dpf, most photoconverted hair cells were only *cabp1b*+, indicating that early-developing hair cells mature to the extrastriolar subtype (Fig 4B).

**Figure 4:**
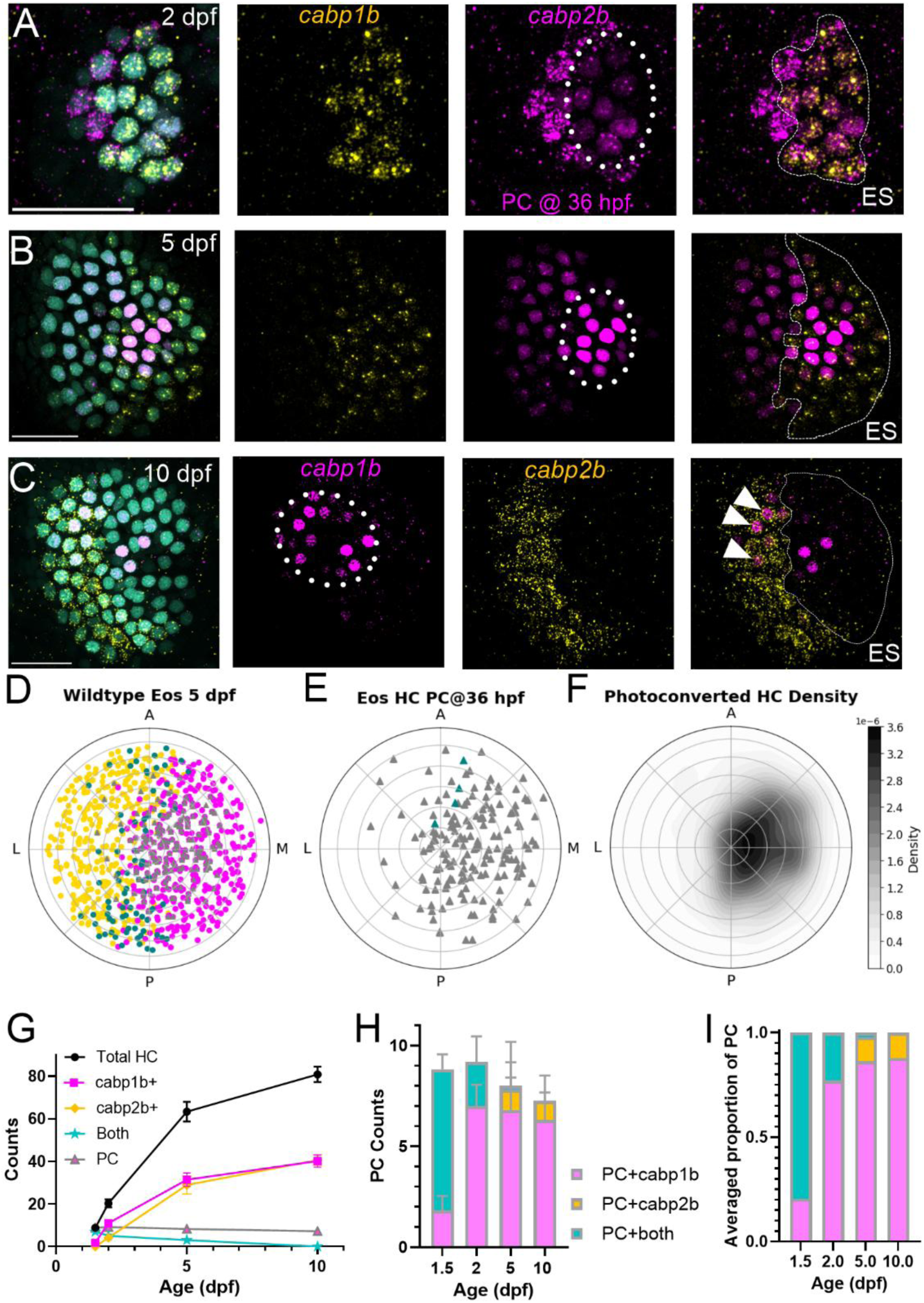
Tracking hair cell fate change during early development. A – C) HCR FISH probing for *cabp1b* and *cabp2b* in *Tg(myo6b:nlsEos)* at 2, 5, and 10 dpf. First-developing hair cells were photoconverted (“PC”) at 36 hpf (dotted circle) to magenta while later-developing hair cells only have unconverted (cyan) Eos. Identity of all hair cells were tested with *capb1b* (“ES”) and *capb2b*. Scale bars = 20 µm. D-F) Polar plot summaries of 5 dpf utricles (B) (n = 11 utricles, 6 fish). PC hair cells (see triangles in E) are mostly located in extrastriola (F). Few PC hair cells were also intermediate (blue triangles in E). G) Summary of developmental data hair cell counts. H-I) Identity of PC hair cells by age (n = 11 ears, 8 fish (2 dpf); 10, 6 (5); 16, 8 (10)). By the end of larval development, most PC hair cells were still extrastriolar (ES), but a few were now striolar (white arrow heads in C)

When we plotted the location of early-developing hair cells (Fig 4D-F), we found they were preferentially located in the extrastriola half of the plot (180° - 360°) (binomial test, 132/163 (81%) vs 50%, p < 0.0001). These observations confirm that early-developing hair cells, regardless of identity, are more likely located in the medial half of the utricle. When we examined the identity of early-developing hair cells at 10 dpf, we found they no longer displayed an intermediate phenotype. We observed that some photoconverted hair cells expressed *cabp2b*+ (Fig 4C-F) at this stage. We saw no significant difference in the number of converted cells over time, suggesting there is little or no loss of cells by turnover, consistent with our previous results (Beaulieu et al., 2024).

Taken together, our results demonstrate that early-developing hair cells have an intermediate phenotype, with later-developing hair cells specified as striolar or extrastriolar. Most early developing cells become incorporated into the medial zone and acquire an extrastriolar phenotype. Hair cells along the midline of the utricle between the two zones are the most likely to retain intermediate phenotype. These results suggest that hair cell differentiation may respond to signaling that indicates spatial localization.

### Retinoic acid synthesizing and degrading enzyme spatially colocalize with distinct zones

In mammalian vestibular end organs, a retinoic acid (RA) gradient drives the differential specification of striolar and extrastriolar hair cells (Ono et al., 2020a, 2020b). We therefore investigated whether the RA synthesis enzyme Aldh1a3 and degradation enzyme Cyp26b1 are expressed in the developing fish ear. Single cell RNA sequencing data indicate that distinct populations of supporting cells differentially expressed genes for *aldh1a3* or *cyp26b1* in the zebrafish maculae (Fig S1F-G). We used HCR FISH to validate spatial expression patterns (Fig 5). At 5 dpf, we observed complementary patterns of *aldh1a3* and *cyp26b1* in and around the utricle (Fig 5A; n = 12 ears, 8 fish). Expression of *aldh1a3* was found in both hair cells and supporting cells in the extrastriolar (medial) half of the utricle, overlapping expression with *cabp1b*+ hair cells (Fig 5B; n = 5 ears, 3 fish). Expression of *cyp26b1* was found in supporting cells in the striolar (lateral) side of the utricle, and colocalized with *cabp2b+* hair cells (Fig 5C; n = 5 ears, 4 fish).

**Figure 5:**
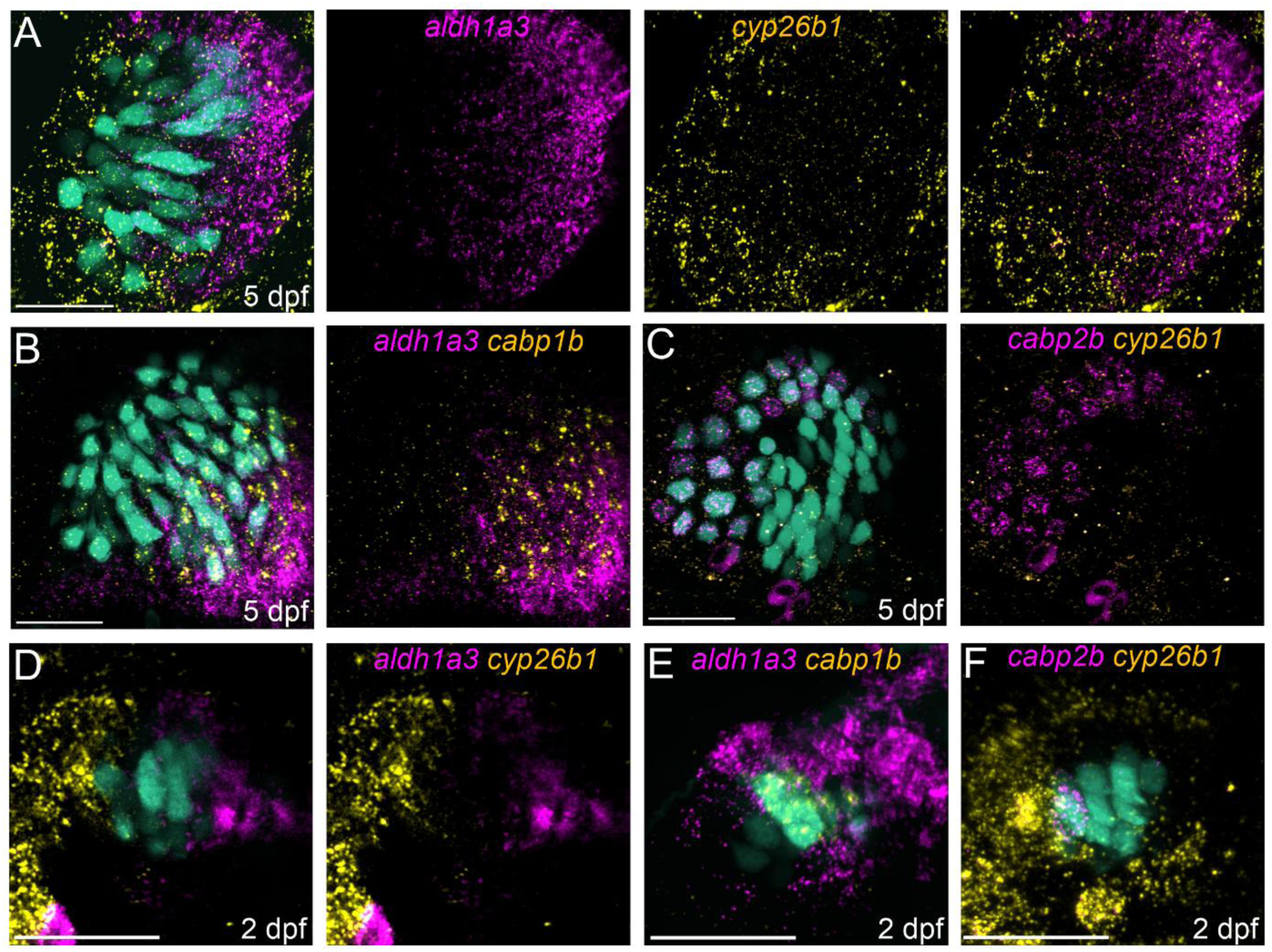
Patterning of retinoic acid synthesizing and degrading enzymes correlate with utricular zones. A) HCR FISH probing for *aldh1a3* (magenta) and *cyp26b1* (yellow) shows complementary patterning of retinoic acid (RA) enzyme in 5 dpf *Tg(myo6b:GFP)* utricle. Scale bars = 20 µm. B) *aldh1a3* (magenta) and *cabp1b* (yellow) indicates colocalization of RA synthesizing enzyme and extrastriolar hair cells. C) *cabp2b* (magenta) and *cyp26b1* (yellow) indicates colocalization of RA degrading enzyme and striolar hair cells. D) At 2 dpf, *aldh1a3* (magenta) and *cyp26b1* (yellow) are polarized to either side of the developing maculae, where a few central hair cells contact neither end. E-F) 2 dpf hair cells expressing *cabp1b* contact *aldh1a3*, as hair cells expressing *cabp2b* contact *cyp26b1*.

We also examined the expression of RA-regulating enzymes at different stages of development (Fig S3). At 1 dpf, when early-developing hair cells have an intermediate identity, we observed that the supporting cells surrounding the nascent hair cells are entirely labeled by *aldh1a3*; *cyp26b1* expression is limited to the otic tissues ventral to the developing maculae (Fig S3A; n = 4 ears, 4 fish). By 2 dpf, the expression patterns are polarized mediolaterally (Fig 5D; n = 9 ears, 5 fish), comparable to that at 5 dpf. *Cyp26b1*-labeled cells contact *cabp2b*+ hair cells on the lateral side of the epithelium (n = 5 ears, 3 fish) and *aldh1a3* labeled cells contact medial *cabp1b*+ hair cells (n = 4 ears, 4 fish) (Fig S3B, Fig 5E-F), leaving a region in the middle of the epithelium where neither enzyme is expressed (Fig 5D). We suggest that this RA-neutral area colocalizes with intermediate hair cells in the utricle at 2 dpf (Fig 3C). At 21 dpf, the lateral extrastriola begins to develop (Bang et al., 2001; Beaulieu et al., 2024), so that the striola eventually becomes surrounded by the extrastriolar region in the adult. At this stage we observed expression of *aldh1a3* on the lateral edge of the utricle (Fig S3C), coinciding with the expansion of the extrastriolar zone to both sides of the periphery (n = 4 ears, 3 fish).

### Treatment with RA alters utricular patterning

We next asked whether increased RA signaling altered zonal patterning during utricular development (Fig 6). We tested various concentrations of RA, dissolved in the fish media while embryos developed between 1 and 5 dpf, after which they were fixed and utricles processed with HCR FISH (Fig 6A-D). Given that RA enrichment promotes the development of the mouse utricular extrastriola, we hypothesized that increasing RA during development would decrease the number of striolar hair cells.

**Figure 6:**
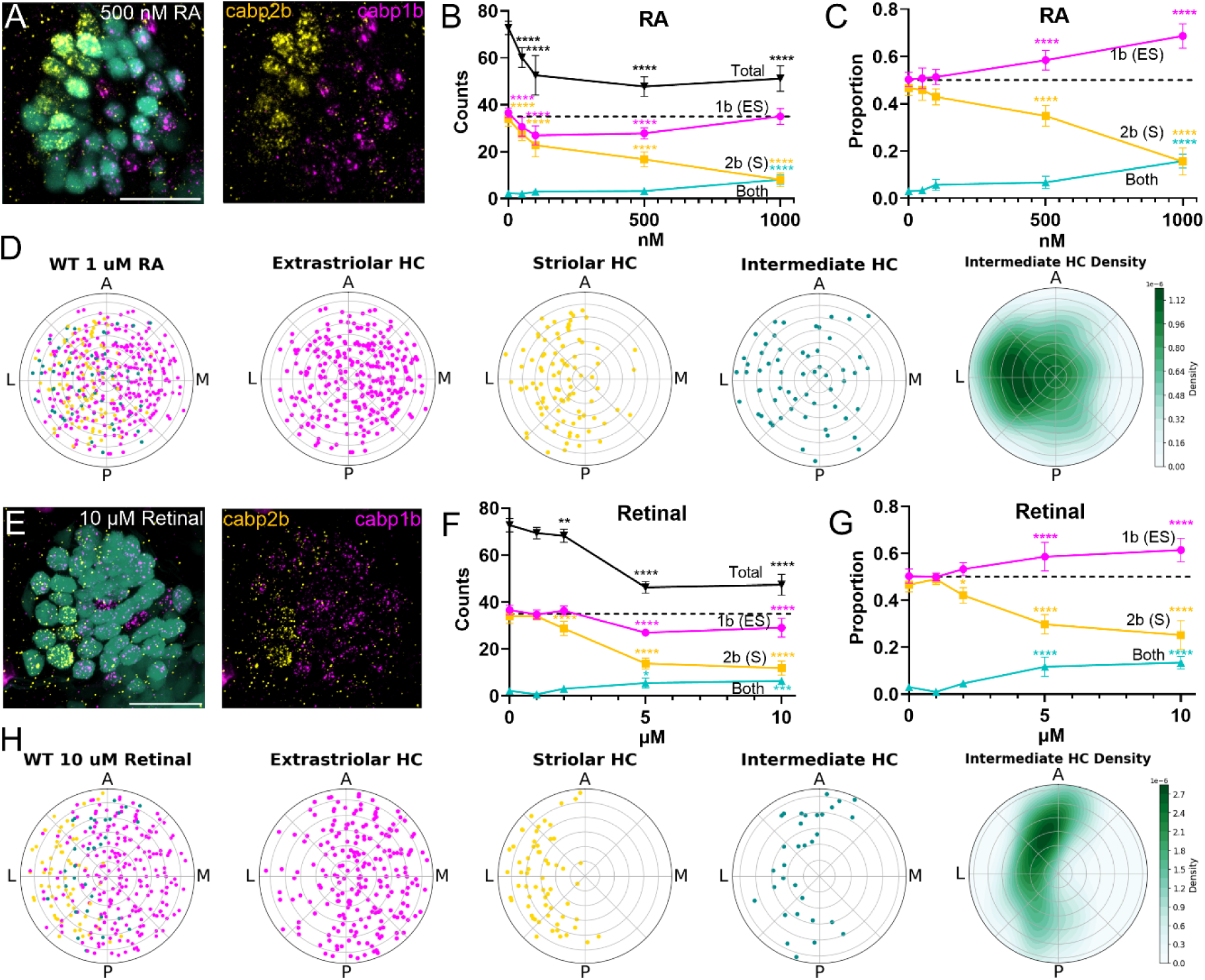
Manipulating retinoic acid altered utricular patterning. A) *cabp1b* (magenta) and *cabp2b* (yellow) expression in *Tg(myo6b:GFP)* utricle at 5 dpf after treatments with 500 nM retinoic acid (RA) for 4 days (1-5 dpf). Scale bars = 20 µm. B-C) Hair cell counts and relative proportions in wildtype fish as a function of RA concentration. Controls: n = 10 ears, 6 fish. RA: n = 11, 11 (50 nM); 12, 11 (100 nM); 15, 11 (500 nM); 8, 6 (1 µM). D) Summary polar plots of utricles after 1 µm RA (n = 8 ears, 6 fish). E) *cabp1b* (magenta) and *cabp2b* (yellow) expression in 5 dpf *Tg(myo6b:GFP)* utricle after treatment with 10 µM retinal. F-G) Hair cell counts and relative proportions in wildtype fish as a function of retinal concentration Retinal: n = 7, 6 (1 µM); 5, 9 (2 µm); 5, 6 (5 µM); 7, 8 (10 µM) H) Polar plot summary of utricles after 10 µM retinal (n = 8, 8).

Relative to non-treated fish, vehicle-treated control wildtype fish had similar proportions of extrastriolar to striolar hair cells (50% extrastriolar and 47% striolar), and few intermediate cells (3%) (Fig 6B-C). At the highest dose (1 µM), we observed the number of extrastriolar hair cells was not significantly different (Two-way ANOVA, 36.6±2.1 vs 35±3.2; p = 0.9), whereas striolar hair cells were reduced (34±3.1 vs 8±3; p<0.0001) and intermediate hair cells increased (2.2±1.0 vs 8.1±2.1; p < 0.0001) (Fig 6B). However, total hair cell counts decreased as the concentration of RA increased, indicating that the utricles had reduced growth (One-way ANOVA, F(4, 51) = 37.3, p < 0.0001) (Fig 6B). At 1 µM treatment, total hair cell number decreased from 72.8 ± 2.9 to 51.1±5.4 (One-way ANOVA, p < 0.0001). To account for this reduction, we compared the relative proportions of hair cell subtypes (Fig 6C). After treatment with 1 µM RA, extrastriolar hair cells made up 70% of the utricle, striolar hair cells 15%, and intermediate hair cells 15%. Thus, we attribute the difference in total hair cells between the control group and experimental groups to a reduction in striolar hair cells.

We also tested the effects of treatment with retinal (Fig 6E-H), which is locally converted to retinoic acid by endogenous Aldh1a3 enzyme. As with RA treatment, we observed a decrease in the total number of hair cells, most significantly at the highest dose (One-way ANOVA, 72.8±2.9 vs 47.4±4.5; p < 0.0001) (Fig 6F). We observed a slight decrease in extrastriolar hair cells (Two-way ANOVA, 36.6±2.1 vs 29.1±4; p < 0.0001), a larger decrease in striolar hair cells (34±3.1 vs 11.9±3.0; p < 0.0001), and an increase in intermediate hair cells (2.2±1 vs 6.4±1.4; p = 0.0006). These changes were reflected in the relative proportions of hair cell types: at the highest concentration, extrastriola = 61%, striola = 25%, intermediate = 14% (Fig 6G).

We plotted the spatial organization of hair cells after RA or retinal treatments in wildtype fish (Fig 6D, H). In wildtype fish, after 1 µM RA, the patterning of striola/extrastriola was significantly disrupted: *cabp1b+* hair cells were significantly more dispersed relative to untreated fish (two-proportion z-test, only 159/265 on the medial side (60%) vs 302/346 (87%), z = -7.7, p < 0.0001) and so were *cabp2b+* hair cells, if to a lesser extent (two-proportion z-test, 80/94 on lateral side (85%) vs 297/332 (89%), z-stat = -1.1, p < 0.0001) (Fig 6D). After 10 µM retinal, *cabp1b+* hair cells were more dispersed relative to untreated utricles (two-proportion z-test, 50/220 (68%), z-stat = -5.5, p < 0.0001), but *cabp2b+* hair cells were more localized in the striola (two-proportion z-test, 60/62 (97%), z-stat = 1.8, p < 0.0001).

We next examined the spatial distribution of intermediate cells after RA or retinal treatment (Fig 6D, H). We found that after treatment with RA intermediate cells are interspersed with striolar cells (binomial test, 39/59 (66%) vs 50%, p < 0.05) (Fig 6D). By contrast, we found that after retinal treatment, intermediate cells remained on the border between striolar and extrastriolar zones even though the striolar zone was reduced (binomial test, 24/32 (75%) vs 50%, p < 0.01) (Fig 6H). Together these results suggest that intermediate cells persist in regions of intermediate RA concentration.

### Loss of mechanotransduction-driven activity alters zonal patterning in *sputnik* mutant fish

We next sought to understand what factors influence the maturation of intermediate cells so that they acquire zonal identity. Hair cells depend on mechanotransduction activity to mature electrical and morphological features (Lui et al., 2016; Corns et al., 2018), stereocilia maturation (Krey et al., 2021), synapse development (Lee et al., 2021) and mitochondrial growth (Mcquate et al., 2023). To test to what extent hair cell activity drives subtype specification in the zebrafish utricle, we assessed zonal organization in 5 dpf *cadherin23^-/-^*(“*sputnik*”) mutant fish (Fig 7), which lack the tip-links necessary to open mechanotransduction channels in hair cells and are deaf and gravity-blind as a result (Söllner et al., 2004).

**Figure 7:**
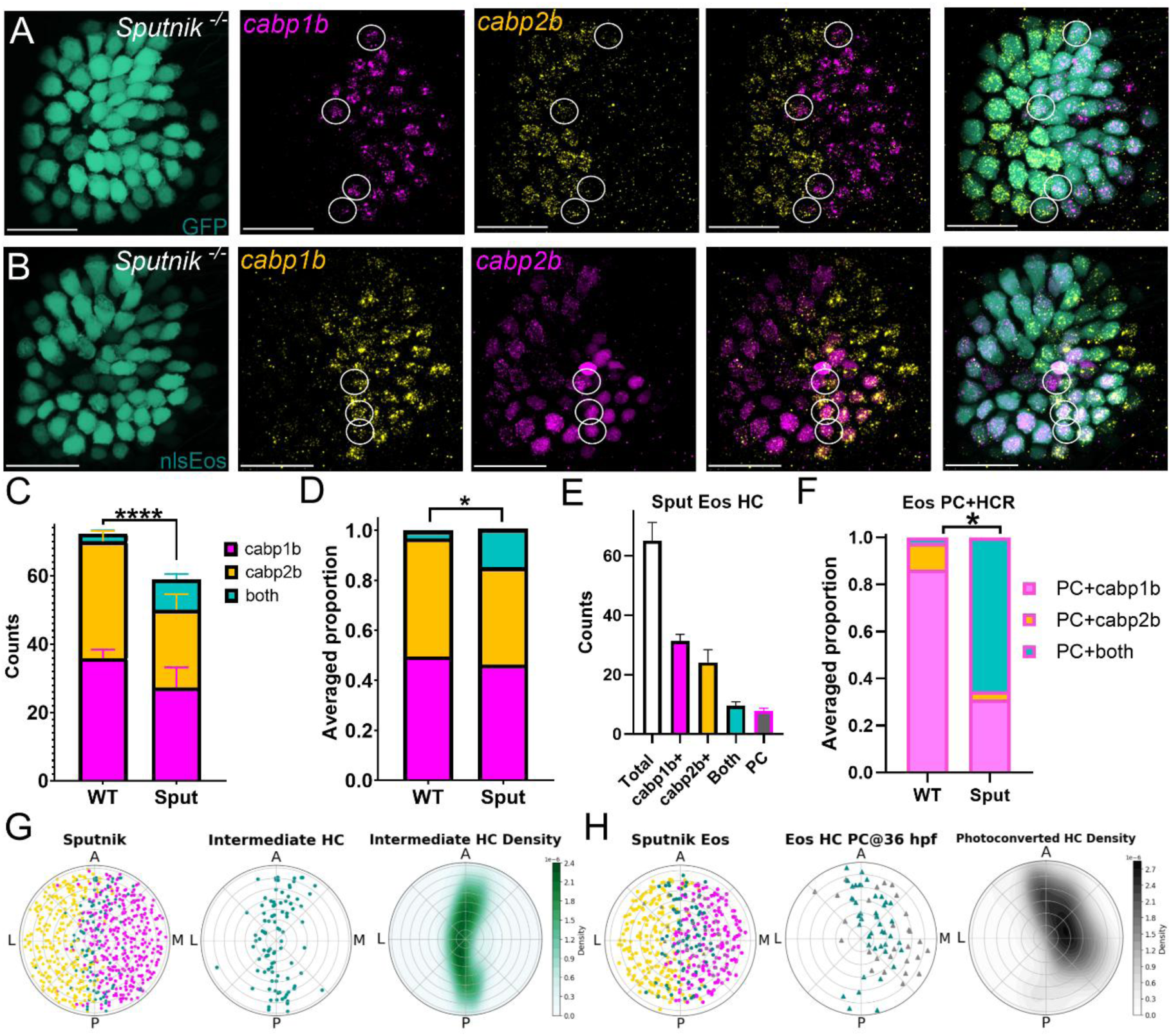
Sputnik mutant fish exhibit abnormal patterning. A) Expression of *cabp1b* and *cabp2b* in *Tg(myo6b:GFP); sputnik* mutant fish at 5 dpf. White circles indicate many intermediate (double-labeled) hair cells present even at 5 dpf. B) HCR FISH after photoconversion at 36 hpf in *Tg(myo6b:nlsEos)* at 5 dpf. White circle indicates hair cell that is not photoconverted but still double-labeled. Scale bar = 20 µm. C-D) *Sputnik* mutants (“Sput”) (n = 21 ears, 15 fish) have fewer hair cells and significantly different proportions of striolar/extrastriolar/intermediate hair cells at 5 dpf relative to wildtype (“WT”) fish (n = 12, 8). E-F) Hair cells photoconverted at 36 hpf in *sputnik* fish with *Tg(myo6:nlsEos)* (n = 7, 7) show that most early-developing hair cells stay in the intermediate state (vs wildtype, data from Fig 4). However, there are fewer photoconverted hair cells than double-labeled hair cells, indicating some later-developing hair cells are intermediate. G) Polar plot summary of 5 dpf utricles from G) *Tg(myo6b:GFP)* (n =12, 9). Intermediate hair cells were located along the zonal boundary. H) Polar plot summary of *Tg(myo6b:nlsEos)* (n = 7, 6). PC hair cells (triangles) are mostly located in extrastriola. Most PC hair cells were also intermediate (blue triangles) and located along the zonal boundary.

Homozygous *sputnik* mutants had significantly fewer utricular hair cells when compared to wildtype fish (Fig 7C) (Welch’s t-test, 72.3±3.1 vs 61.5±9.0, p <0.0001, effect size = 1.4). Therefore, we normalized cell counts to proportions to assess differences in cell type distribution (Fig 7D). While there is a similar proportion of extrastriolar (*capb1b*+) hair cells (Two-way ANOVA, 0.49±0.03 vs 0.46±0.04; p = 0.33, power = 0.06), there is a decrease of striolar hair cells in *sputnik* mutants (0.47±0.03 vs 0.39±0.1; p < 0.0001, effect size = 0.8). We also observed a five-fold increase in the fraction of intermediate-type hair cells present at 5 dpf (Two-way ANOVA, 0.03±0.01 vs 0.15±0.02; p<0.0001; effect size = 4.8) (Fig 7C-D). In all, the distribution of hair cells type in *sputnik* mutants was different relative to wildtype fish (Chi-square test, χ^2^ = 6.6, p < 0.05, effect size = 0.58).

We also assessed zonal distribution in *ca_V_1.3a^-/-^*(“*gemini*”) mutant fish, which lack the calcium channels necessary for neurotransmitter release from hair cells and are also deaf and gravity-blind (Fig S4). Total hair cell counts, *cabp1b*+, *cabp2b*+, and persisting intermediate hair cells were comparable between *gemini* mutants and wildtype fish (Chi-square test, χ^2^ = 0.02, p = 0.99, power = 0.01). This suggests that mechanotransduction driven activity is specifically necessary to drive hair cell zonal maturation.

We next determined the spatial organization of 5 dpf *sputnik* mutant utricles (Fig 7G-H). We observed that, out of 745 hair cells, 47% were *cabp1b*+, 39% were *cabp2b*+, and 14% were intermediate. Like in wildtype fish, *cabp1b+* and *cabp2b+* cells neatly align with the extrastriolar (binomial test, 317/ 354 (90%) vs 50%, p < 0.0001) and striolar zones, respectively (273/299 (95%), p < 0.0001). The increased number of the intermediate hair cells also were along the striolar/extrastriolar boundary (Fig 7G): 69 out of 103 cells (67%) were in this quarter section of the plot (vs 25%, p < 0.0001).

To test whether intermediate-type hair cells in *sputnik* mutants at 5 dpf are simply early-developing hair cells that persisted in an immature state, we crossed *Tg(myo6b:nls-Eos)* with the *sputnik* line to track early-developing hair cells in mutant fish (Fig 7B, E-F). We photoconverted embryos at 36 hpf, fixed them at 5 dpf, and assessed hair cell identity in mutant fish with HCR FISH (Fig 7B, E). Indeed, most photoconverted hair cells were intermediate (*cabp1b* + *cabp2b*) (∼60%) (Fig 7F). Some, however, matured to become extrastriolar (∼35%) and only a very few became striolar (∼5%) (Chi-square test, χ^2^ = 8.75, p = 0.01, effect size = 1) (Fig 7F). We plotted the 5 dpf spatial organization of early-developing hair cells (Fig 7H). These were preferentially located in the extrastriola half of the plot (180° - 360°) (binomial test, 52/72 (73%) vs 50%, p < 0.001). We also noted that early-developing cells with persistent intermediate phenotypes were correspondingly in the extrastriola (24/29 (83%) vs 50%, p < 0.001). Intriguingly, some late-developing hair cells were also intermediate (Fig 7B), suggesting that hair cell mechanotransduction activity continues to play a role in driving hair cell identity after early-developing hair cells have matured.

We noted that both loss of mechanotransduction and increase in RA signaling result in similar zonal patterning defects: a decrease in the striolar zone and increase in intermediate hair cells. However, when probing for *aldh1a3* and *cyp26b1* using HCR FISH, we observed that *sputnik* (n = 11 ears, 7 fish) and *gemini* (n = 16, 11 fish) mutant fish exhibited normal *aldh1a3*/*cyp26b1* complementary patterning (Fig S5A-B). We also observed normal complementary patterning of *aldh1a3*/*cyp26b1* in *atoh1a* mutant fish (Fig S5C; n = 9 ears, 7 fish), which lack transcription factor Atoh1A necessary for hair cell differentiation. Together these results suggest that patterned expression of RA signaling enzymes is independent of hair cells.

We next tested the effect of RA or retinal treatment on utricular patterning in *sputnik* mutant fish (Fig 8). After 10 µM retinal or 1 µM RA, just like treated wildtype fish, *sputnik* mutant fish showed an overall decrease in the number of hair cells relative to vehicle-treated mutants and wildtypes (Fig 8B) (One-way ANOVA, p<0.0001). After 10 µM retinal, the average number of extrastriolar hair cells remained unchanged in retinal-treated *sputnik* fish relative to vehicle-treated mutants (Two-way ANOVA, 31.8±4.7 vs 31.7±4; p = 1) or retinal-treated wildtypes (31.8±4.7 vs 29.1±4.0, p=0.9) (Fig 8B). The number of intermediate type cells was also not different relative to vehicle-treated mutants (Two-way ANOVA, 9.8±3.4 vs 8.5±2.1; p=1) or retinal-treated wildtypes (9.8±3.4 vs 6.4±1.4; p = 0.3) (Fig 8B). However, striolar hair cells significantly decreased relative to mutant controls (Two-way ANOVA, 5.4±1.9 vs 21.7±3.8; p<0.0001) and to retinal-treated wildtypes (5.4±1.9 vs 11.9±3.0; p<0.0001). RA-treated *sputnik* fish also showed a significant decrease in the number of striolar hair cells relative to vehicle-treated mutants (Two-way ANOVA, 10.7 ± 2.2 vs 21.7±3.8; p<0.0001), a slight decrease in extrastriolar hair cells (26.9±7.2 vs 31.4±4.1; p = 0.16), but no significant change in intermediate (10.7±1.2 vs 8.5±2.1; p = 1) (Fig 8B).

**Figure 8:**
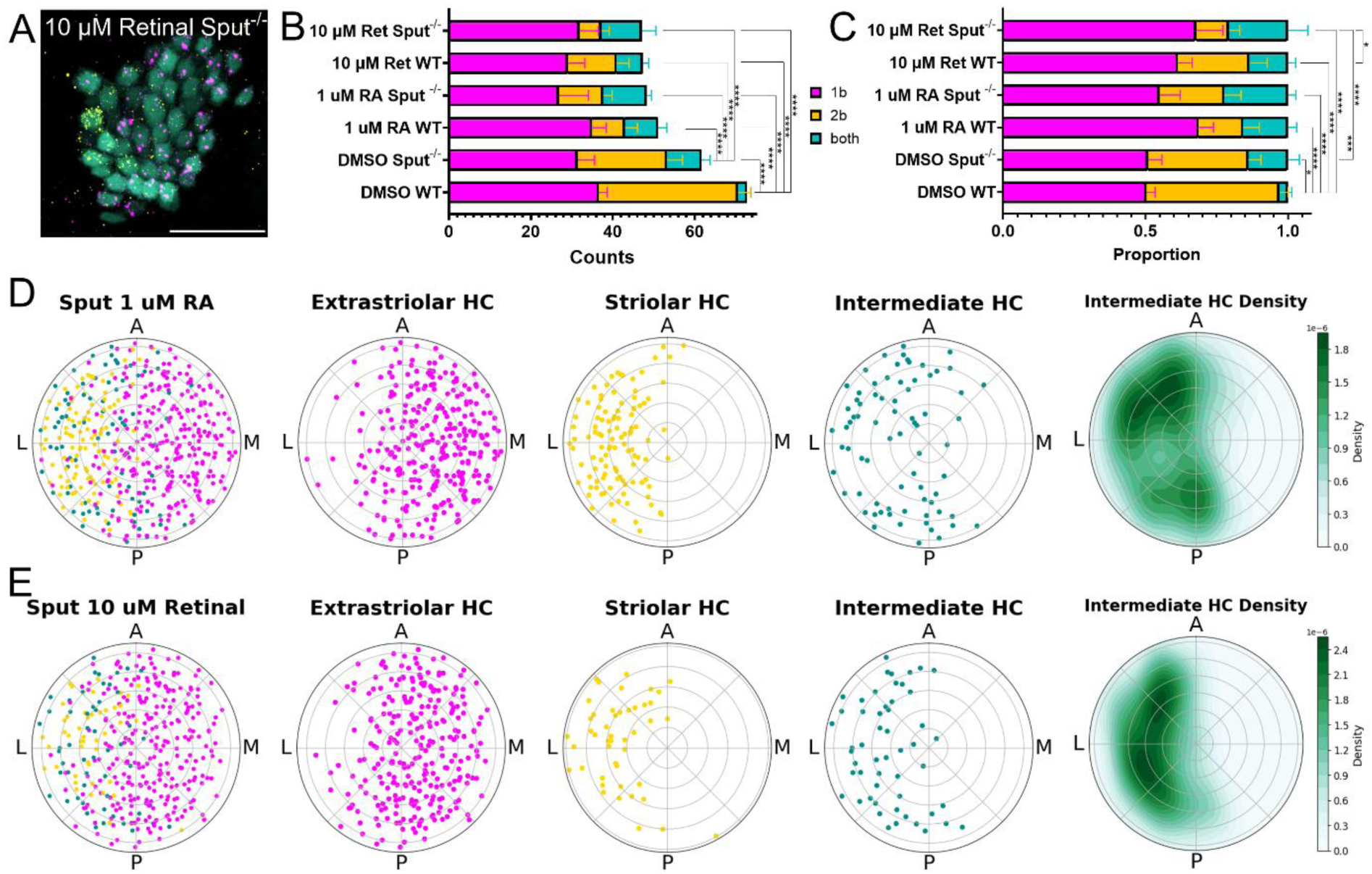
RA further disrupted patterning in *sputnik* fish. A) *cabp1b* (magenta) and *cabp2b* (yellow) expression changes in *sputnik Tg(myo6b:GFP)* utricle after 10 uM retinal treatment (scale bar = 20 µm). B) *Sputnik* mutant fish treated with either 1 µM RA (n = 9, 6) or 10 µM retinal (n= 7, 7) show a decrease in the overall number of hair cells relative to sputnik DMSO controls (n = 17, 9) or WT DMSO controls. C) Relative distribution of hair cells changed in RA and retinal-treated WT and *sputnik* mutant fish relative to controls. D-E) Summary polar plots of *sputnik* mutant utricles after D) 1 µm RA (n = 10, 6) or E) 10 µM retinal (n = 8, 8) show a disruption of striolar/extrastriolar patterning and an increase in intermediate hair cells in the reduced striolar zone.

The hair cell type distribution in retinal-treated *sputnik* mutants was different relative to retinal-treated wildtype fish (Fig 8C) (Chi-square test, χ^2^ = 7.6, p = 0.02, effect size = 0.3), vehicle-treated *sputniks* (χ^2^ = 16.3, p < 0.000, effect size = 0.5), and vehicle-treated wildtype fish (χ^2^ = 38.6, p < 0.000, effect size = 1.2) (Fig 8C). Thus, *sputnik* mutant fish show an additional loss of the striolar zone in response to increased retinoic acid.

We plotted the spatial organization of hair cells after RA or retinal treatments in *sputnik* fish (Fig 8D-E). RA- or retinal-treated *sputnik* mutant fish demonstrated pattern disruption: after 1 µM RA, *cabp1b+* hair cells were significantly dispersed relative to untreated WT fish (two-proportion z-test, 198/281 (70%) vs 87%, z-stat = -5.2, p = 0), *cabp2b+* hair cells were more localized in the striola (two-proportion z-test, 106/109 (97%) vs 89%, z-stat = 2.5, p = 0), and intermediate hair cells localized in the striola (binomial test, 66/83 (80%) vs 50%, p = 0) (Fig 8D). After 10 µM retinal, *cabp1b+* hair cells were more distributed (two-proportion z-test, 189/272 (69%) vs 87%, z-stat = -5.4, p = 0), the *capb2b+* hair cells more limited (two-proportion z-test, 42/46 (91%) vs 89%, z-stat = 0.4, p = 0.07), and the intermediate cells were concentrated in the striolar zone (binomial test, 46/53 (87%) vs 50%, p = 0) (Fig 8E). This further demonstrates that application of retinoic acid or retinal during utricular growth is detrimental to development of the striolar zone.

In summary, we proposed that hair cell zonal patterning in the utricle is dependent upon both mechanotransduction-driven activity and a RA gradient. We compared spatial patterns of extrastriolar, striolar and intermediate cells across all conditions as kernel density estimation (KDE) plots (Fig 9). We also performed spatial autocorrelation analysis of hair cell distributions comparing nearest neighbors for a statistical estimation of their similarities (Fig S6). During development, the RA gradient forms across the lateral-medial axis, driving distinct striolar and extrastriolar hair cell phenotypes (Fig 9A). The intermediate phenotype persists at the striolar/extrastriolar boundary, where retinoic acid is neither enriched nor deficient. Disrupting the RA gradient by increasing RA results in the reduction of the striola and increases in the area occupied by intermediate types (Fig 9A’), resulting in significant differences in spatial autocorrelations compared to wildtype control (Fig S6A). Addition of retinal, which requires endogenous Aldh1a3 to be locally converted to RA, also resulted in loss of striola, but intermediate cells remained at the zonal border (Fig 9A’’), and significant differences in spatial autocorrelation (Fig S6A). The relative proportion of intermediate hair cells increased in *sputnik* mutant fish, which have mechanotransduction dysfunction (Fig 9B), with little change in spatial correlation (Fig S6A). Increasing RA in *sputnik* either directly through RA application or indirectly through retinal exposure exacerbated the disruption in patterning, resulting an increased loss of the striolar zone and increased number of intermediate hair cells (Fig 9B’, B’’) with similar changes in spatial autocorrelation between treatments (Fig S6B).

**Figure 9:**
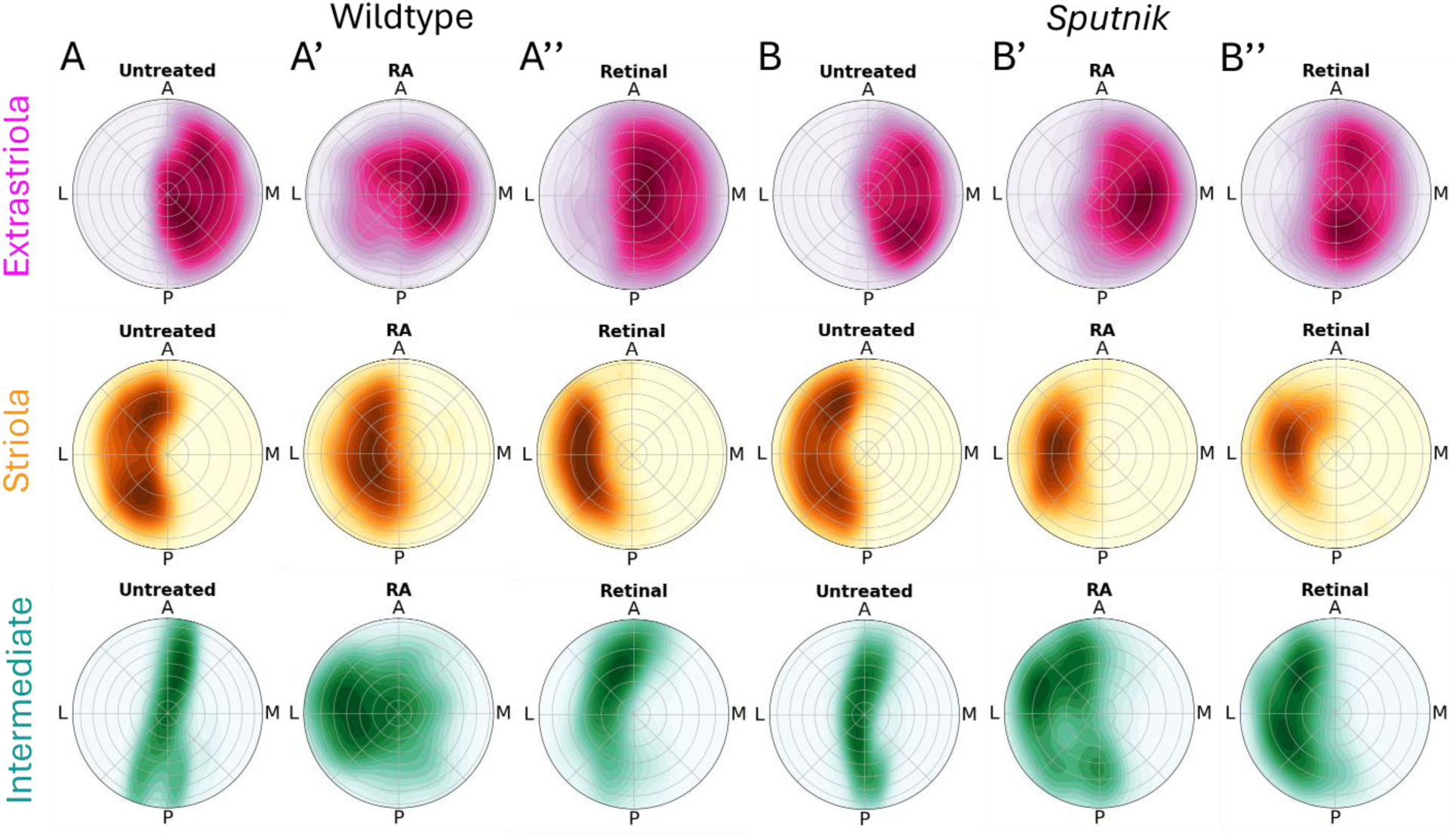
Normal zonal patterning of hair cells depends on RA gradient and activity. Summary KDE plots of polar plots of utricles from Figures 4, 6, 7 and 8. (A) wildtype and (B) *sputnik* fish, at 5dpf (untreated), after 1 µM RA (A’, B’), or 10 µM retinal (A’’, B’’) treatment.

## Discussion

In this study, we took advantage of transcriptomic markers that distinguish different cell subtypes at very early embryonic stages to clarify the dynamics of hair cell specification in the developing zebrafish inner ear. We identified that, in the zebrafish, early-developing hair cells are initially “intermediate” hair cells that later mature into striolar or extrastriolar types; most later-developing hair cells are specified to either type, depending on the zone they develop in. Cell fate tracing demonstrated that the vast majority of early-developing cells mature into extrastriolar hair cells. By contrast, Lui et al., (2022) described striolar hair cells as more mature than extrastriolar cells based on morphological features (long kinocilium, many synaptic ribbons, and innervation by myelinated afferents). We suggest that these morphological features may reflect differences in function across zones rather than maturity.

### Asymmetry of otolith organ development

In zebrafish, each sensory epithelium of the inner ear follows a different developmental timeline. The utricle and the saccule are the first to develop around 1 dpf (Haddon and Lewis, 1996; Riley et al., 1997; Haddon et al., 1999; reviewed in Whitfield et al., 2002; Tanimoto et al., 2011). The cristae begin to form by 2.5 dpf, although these may not be functional until the fish is big enough to induce sufficiently large inertial forces to induce endolymph flow (∼30 dpf) (Bever and Fekete, 2002; Beck et al., 2004, Lambert et al., 2008). The lagena then emerges by the third week, followed by the macula neglecta (Haddon and Lewis, 1996; Bang et al., 2001; Bever and Fekete, 2002; Higgs et al., 2003). After the embryonic stage, in each epithelium hair cells are added continuously so that they are sized in proportion to the fish’s body length (Higgs et al., 2002; Beaulieu et al., 2024).

Although it is present from early developmental stages, we did not rigorously assess the saccule because of the challenges of imaging the posterior epithelium, particularly at older timepoints: its location deep in the ear, and its ninety-degree orientation relative to the utricle made it difficult to image clearly. However, we gathered enough data to appreciate the similarities and differences in the developmental trajectories between the two otolith epithelia, also described in earlier studies (Kwak et al., 2006; Sapède and Pujades, 2010). We noted that in the first five days, the saccule added hair cells at different rates relative to the utricle (Fig 2) (see also Sapède and Pujades, 2010). This may not be completely unexpected as the two epithelia are theorized to serve distinct sensory functions: the utricle is vital for gravity detection and the initial survival of the larvae (Riley and Moorman, 2000; Kwak et al., 2006; Mo et al., 2010), while the saccule is implicated as the primary hearing organ in the larval fish (Lu and DeSmidt, 2013; Inoue et al., 2013; Yao et al., 2016; Lara et al., 2022; Lau and Vasconcelos, 2023).

Evidence also suggests that the development of the utricle and saccule may be independently regulated. In other organisms, the development of the vestibular and hearing organs is shown to be under the control of different regulators (Choo et al., 1998; Riccomagno et al., 2002; Son et al., 2015). In the developing chick ear, for example, the vestibular organs were more susceptible to RA than the hearing organs (Choo et al., 1998). Sapède and Pujades (2010) investigated the role of Hedgehog (Hh) signaling in saccular development in the larval fish given the role of Hh signaling in posterior otic specification (Hammond et al., 2003). They showed that interrupting Hh signaling reduced the number of hair cells in the saccule but not the utricle. Future experiments may examine whether RA plays a role in saccular formation in larval fish.

### Stage-dependent roles for RA signaling during inner ear development and growth

The spatiotemporal regulation of a RA gradient through enzyme mediated synthesis (Aldh1a) and degradation (Cyp26) of RA is fundamental to the formation of the striolar/extrastriolar zones of the otolith organs and the central/peripheral zones of the cristae of the mouse (Ono et al., 2020a, 2020b). We found that the spatial complementary patterning of *aldh1a3* in the extrastriola and *cyp26b1* in the striola is evolutionarily conserved between the zebrafish and the mouse. We hypothesize that this RA gradient across the anterior/posterior axis in the utricle drives hair cell specification in the lateral/medial sides of the epithelia.

Retinoic acid (RA) has multiple functions during inner ear development and maintenance through adulthood (Corey and Breakfield, 1994; Romand et al., 2001, 2006, 2013; Whitfield et al., 2002; Hans and Westerfield, 2007; Frenz et al., 2010; Bok et al., 2010; Ono et al., 2020a, 2020b; Mackowetzky et al., 2022). In the fish, RA signaling is key for normal development of the early otic capsule (Whitfield et al., 1996; Whitfield et al., 2002, Hans and Westerfield, 2007; Mackowetzky et al., 2022). Here we provide evidence for RA signaling in zonal specification of the sensory epithelia at later stages, a function conserved in mouse inner ear patterning. Our initial analysis suggests that a second zone of *aldh1a3* expression appears as the lateral extrastriolar zone begins to develop at about 3 weeks of age, near the end of larval development, resulting in a central shift of the striola. Given that the sensory organs of the zebrafish inner ear undergo continuous growth (e.g. Bang et al., 2001; Higgs et al., 2002), it is intriguing to think that RA signaling may continue to provide spatial information throughout the life of the animal.

### Activity during hair cell development drives maturation

We were struck by the observation that intermediate hair cells had transcriptomic markers for both types and how they were located along the anterior/posterior axis. We expect that most of the intermediate hair cells persist at the boundary between the zones because neither RA enzyme is enriched relative to the other. We suggest that hair cells located in this “RA-neutral” boundary are thus more dependent upon other cues for their acquisition of a specific zonal identity.

We propose that mechanotransduction activity contributes to hair cell maturation. Like other vertebrates, mechanotransduction in the fish ear requires the tip link, a precise chain of proteins that link the stereocilia and serve as the local mechanical force to open transduction channels; breaking this chain decimates hair cell function (Söllner et al., 2004; Kindt et al., 2012; Maeda et al., 2017; Erickson et al., 2020). In *sputnik* mutant fish with disruptions in the *cdh23* gene, where hair cells fail to form tip links and thus have little to no spontaneous activity (Seiler and Nicolson, 1999; Trapani and Nicolson, 2011), the majority of intermediate hair cells never mature. These early-developing hair cells localize at the center of the utricle close to the striolar/extrastriolar boundary, a region where we predict RA signals to potentially be less effective in conferring regional identity.

Hair cells show spontaneous activity prior to hearing onset in mice, with a role in their maturation (Tritsch et al., 2010; Eckrich et al., 2019; De Faveri et al., 2025). Depolarization in immature inner hair cells drives calcium transients that shape hair cell morphology, refine synapses, and enhance intercellular coupling with supporting cells (Tritsch et al., 2010; Eckrich et al., 2019; Kersbergen et al., 2023; Hussain et al., 2024; De Faveri et al., 2025). Mechanotransduction, specifically, is hypothesized to be necessary to change hair cell activity from spiking to graded depolarization (Corns et al., 2018), promote stereocilia growth and patterning (Krey et al., 2020) and mature synaptic connections (Lee et al., 2021). In zebrafish, activity is required for maturation of hair cell mitochondrial networks (McQuate et al., 2023). There is also evidence that hair cell mechanotransduction and synaptic activity drives specification of afferent neuron subtypes the cochlea (Shrestha et al. 2018; Sun et al., 2018). Thus, spontaneous hair cell activity prepares the hair cell itself and promotes the maturation of auditory pathways. The mechanisms by which mechanotransduction activity drives hair cell regional specification are unknown. It is well-established that calcium influx promotes neuronal maturation by activating gene transcription through conserved signaling cascades (Greer and Greenberg, 2008; Hagenston et al., 2020). Mutations to the *Ca_V_1.3* calcium channel gene inhibit hair cell maturation (Brand et al., 2003; Jeng et al., 2020) and in zebrafish (McQuate et al., 2023), however we observed no differences in zonal patterning in our current studies.

## Data availability

The associated code is accessible in the following GitHub repository: https://github.com/raible-lab/utricle_spatial_analyses

The datasets generated and analyzed for both molecular biology and modeling experiments can be found in our Dryad repository: 10.5061/dryad.hmgqnk9xs

