## supplemental figures and legends for "Activity and retinoic acid drive hair cell spatial patterning in the zebrafish utricle"

**Supplemental Figure 1**

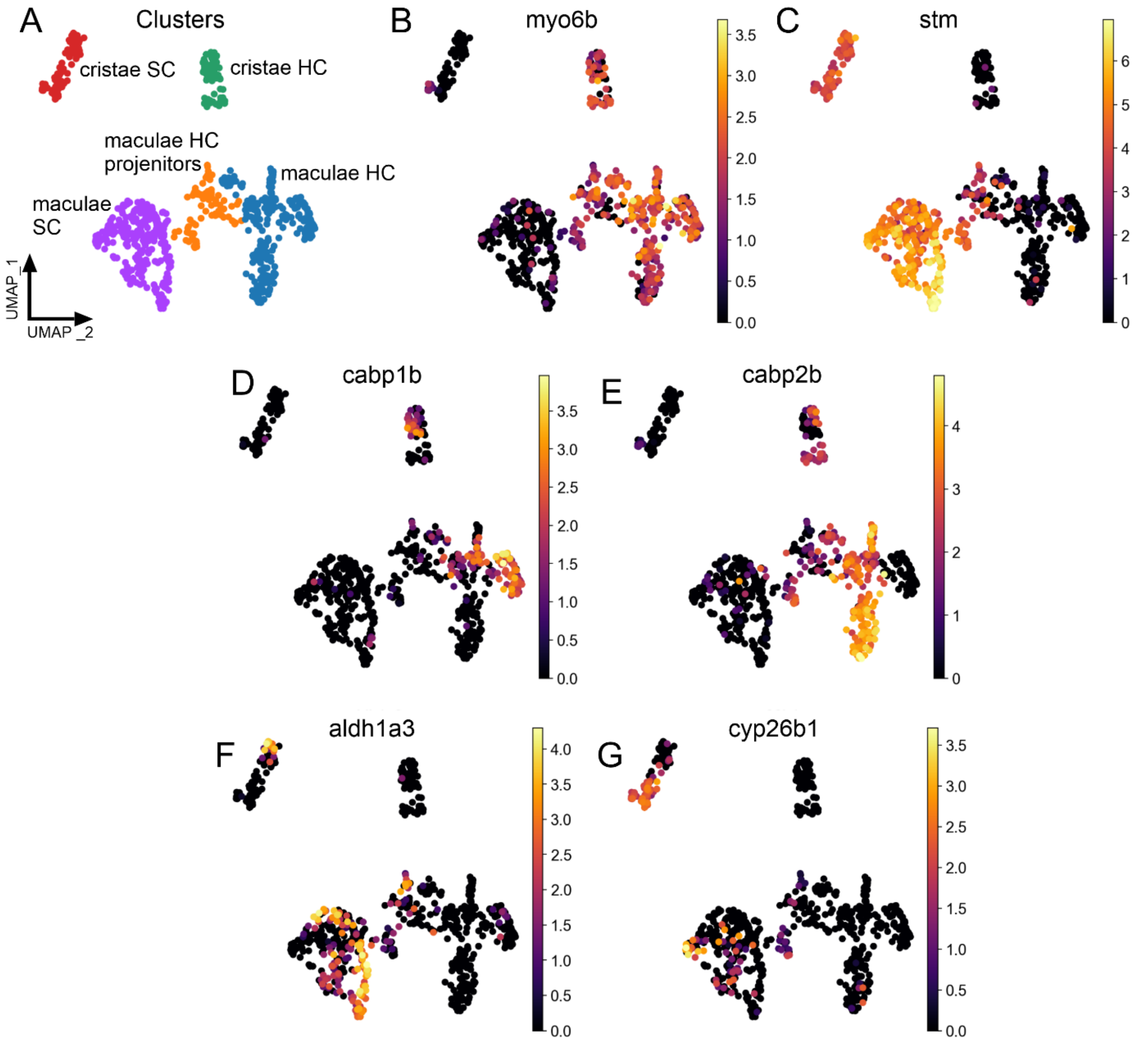


**Figure S1 Single cell RNA sequencing data of larval zebrafish inner ear hair cells and support cells.**

(A) UMAP projection of cell grouped by cell type using scRNA seq data from Daniocell. HC: hair cells, SC: support cells.

(B-C) Hair cells were identified using known marker (B) *myo6b* and support cells by (C) *stm*.

(D-E) Hair cells in the maculae can be differentially identified by (D) *cabp1b* or (E) *capb2b*.

(G-H) Supporting cells show differential expression of retinoic acid synthesizing (G) or degrading (H) enzyme RNA.

**Supplemental Figure 2**

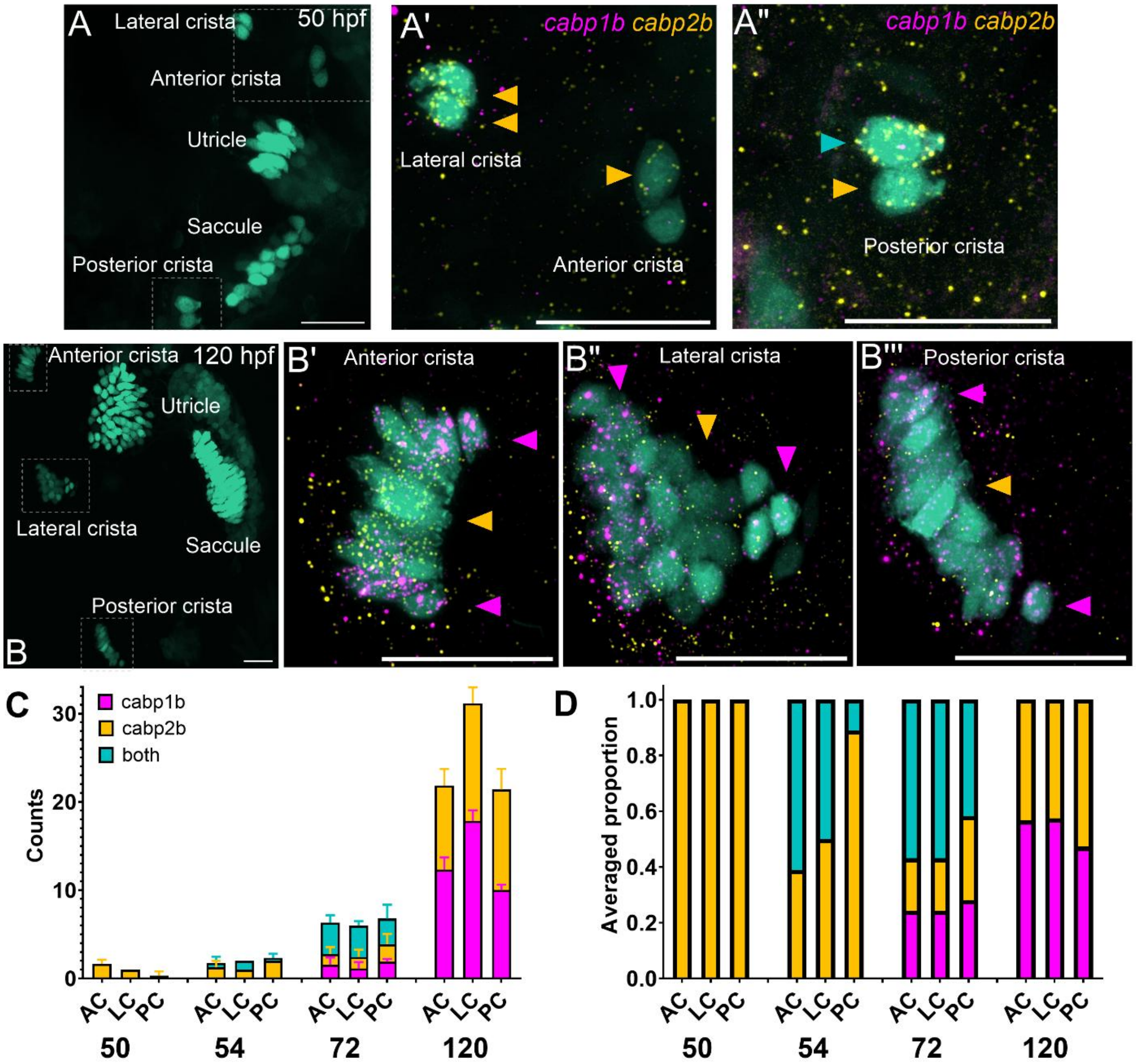


**Figure S2: Identification of cristae hair cell types during larval development.**

A-B) Maximum projections of dorsal views of the inner ear with HCR FISH probing for *cabp1b* and *cabp2b* in the cristae of *Tg(myo6b:GFP)* at (A) 50 and (B) 120 hpf. Scale bar = 20 µm.

C-D) Hair cells are added in the first 5 days in the cristae at different rates and the relative proportions of hair cell subtypes change during larval development (n = 3 (50 hpf); 3 (54); 9 (72); 6 (120).

**Supplemental Figure 3**

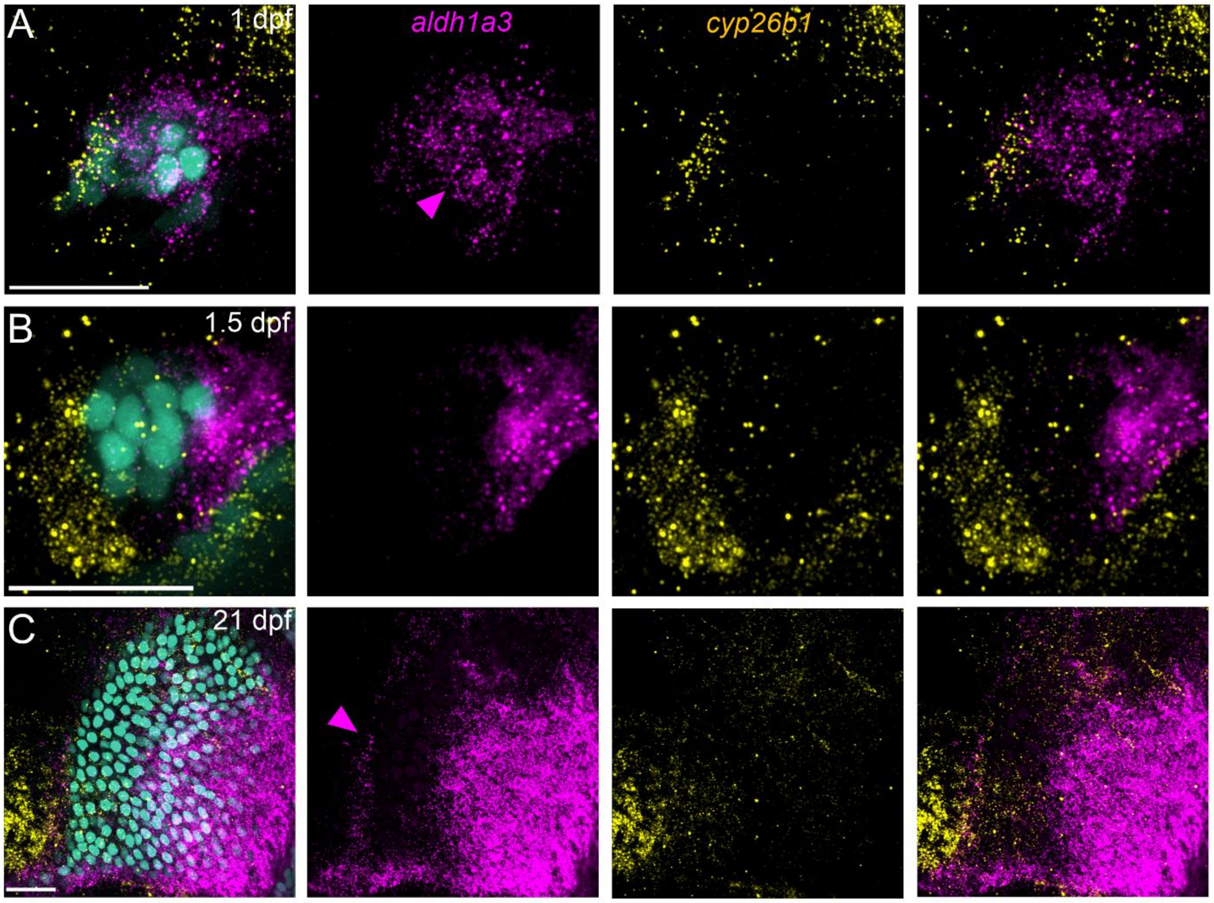


**Figure S3: Patterning of RA enzymes changes with development**

A) Maximum projection of a *Tg(myo6b:GFP)* utricle at 1 dpf with probing for *aldh1a3* (magenta) and *cyp26b1* (yellow) shows that very early hair cells are surrounded by *aldh1a3* (magenta arrowhead). Scale bar = 20 μm.

B) At 1.5 dpf (36 hpf), *aldh1a3* has polarized medially as *cyp26b1* expression develops laterally.

C) By 21 dpf, *aldh1a3* expression begins to develop laterally to *cyp26b1* (magenta arrowhead) as the lateral extrastriola begins to form as exemplified here in a *Tg(myo6b:nlsEos)* utricle.

**Supplemental Figure 4**

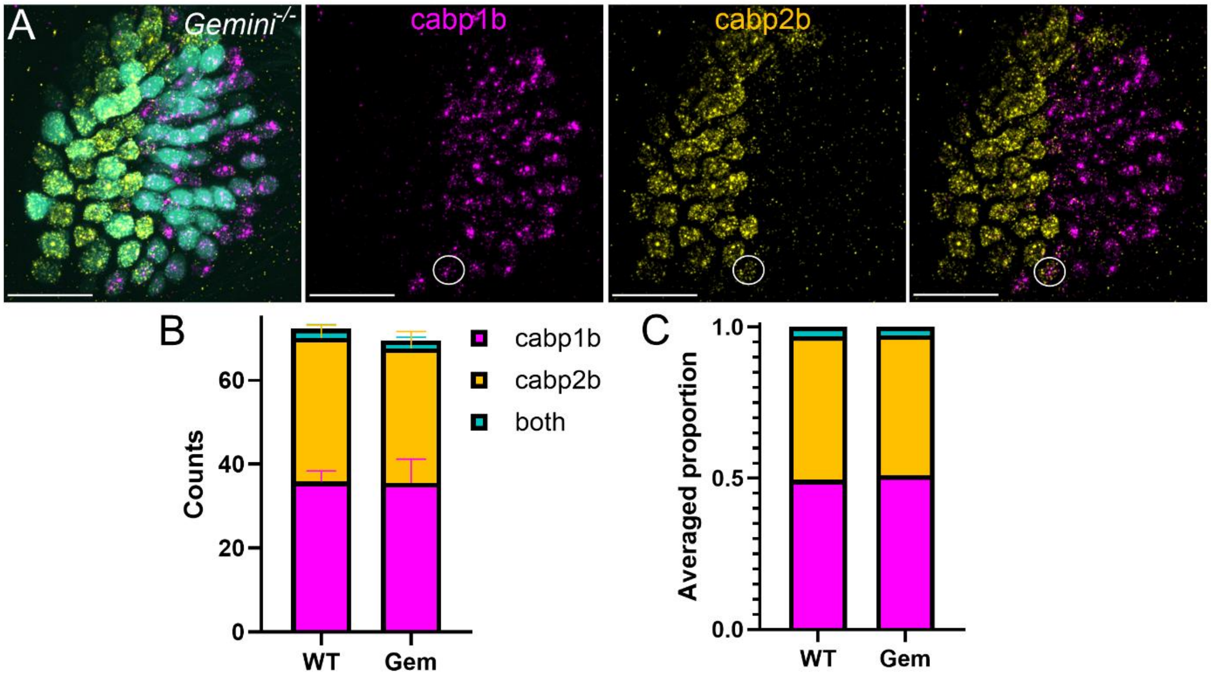


**Figure S4: *Gemini* mutant fish have normal utricular patterning.**

A-B) Maximum projection of *Tg(myo6b:GFP*) utricles in *gemini* mutant fish with HCR FISH probing for *cabp1b* and *cabp2b*. White circle indicates few intermediate (double-labeled) hair cells present at 5 dpf. Scale bar = 20 µm.

C-D) *Gemini* mutants ("Gem") have a similar number of hair cells and similar proportions of striolar/extrastriolar/intermediate hair cells relative to wildtypes at 5 dpf.

**Supplemental Figure 5**

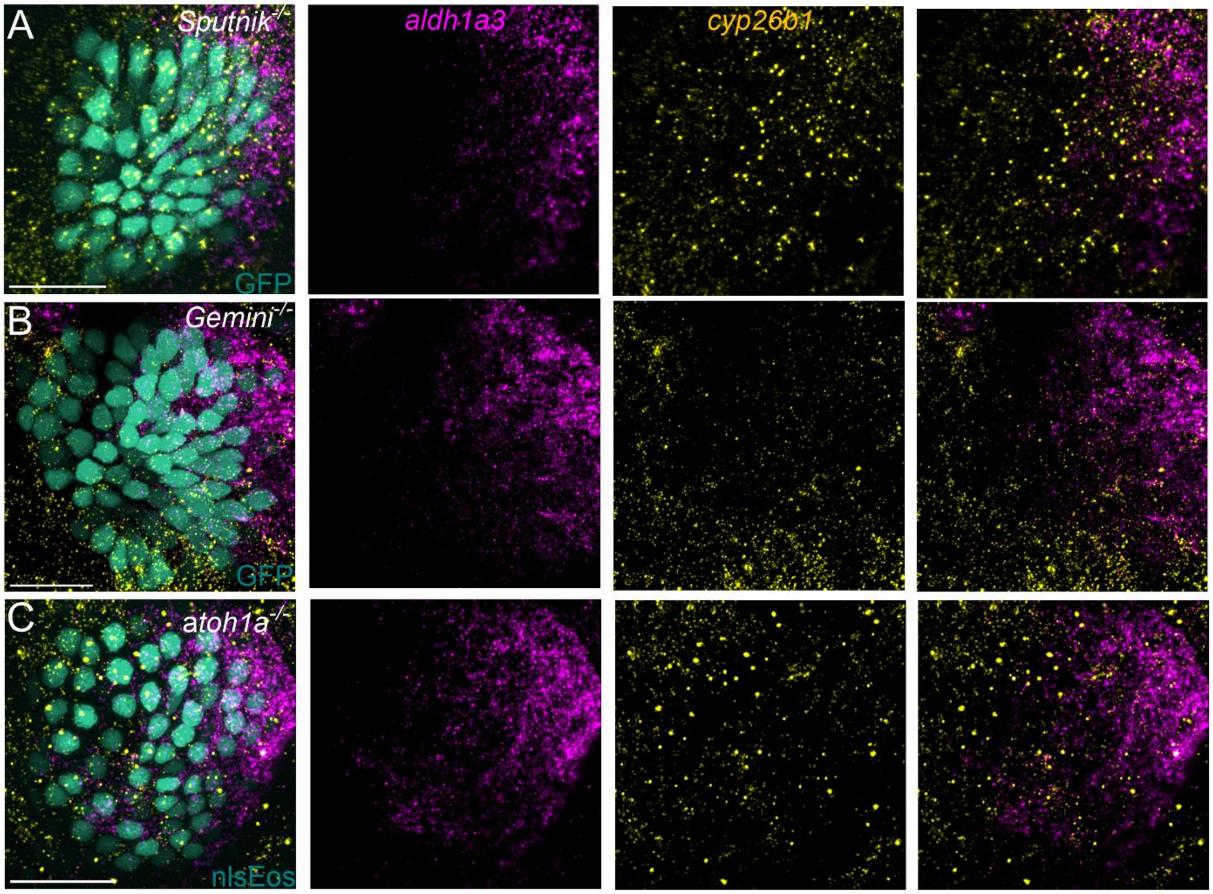


**Figure S5: Fish with hair cell mutations exhibit normal RA enzyme patterning.**

Maximum projections of 5 dpf *Tg(myo6b:GFP)* or *Tg(myo6b:nlsEos)* utricles. Scale bar = 20 µm. HCR FISH probing for *aldh1a3* (magenta) and *cyp26b1* (yellow) shows complementary patterning of retinoic acid (RA) enzyme in A) *sputnik*, B) *gemini*, and C) *atoh1a* mutants are comparable to wildtype.

**Supplemental Figure 6**

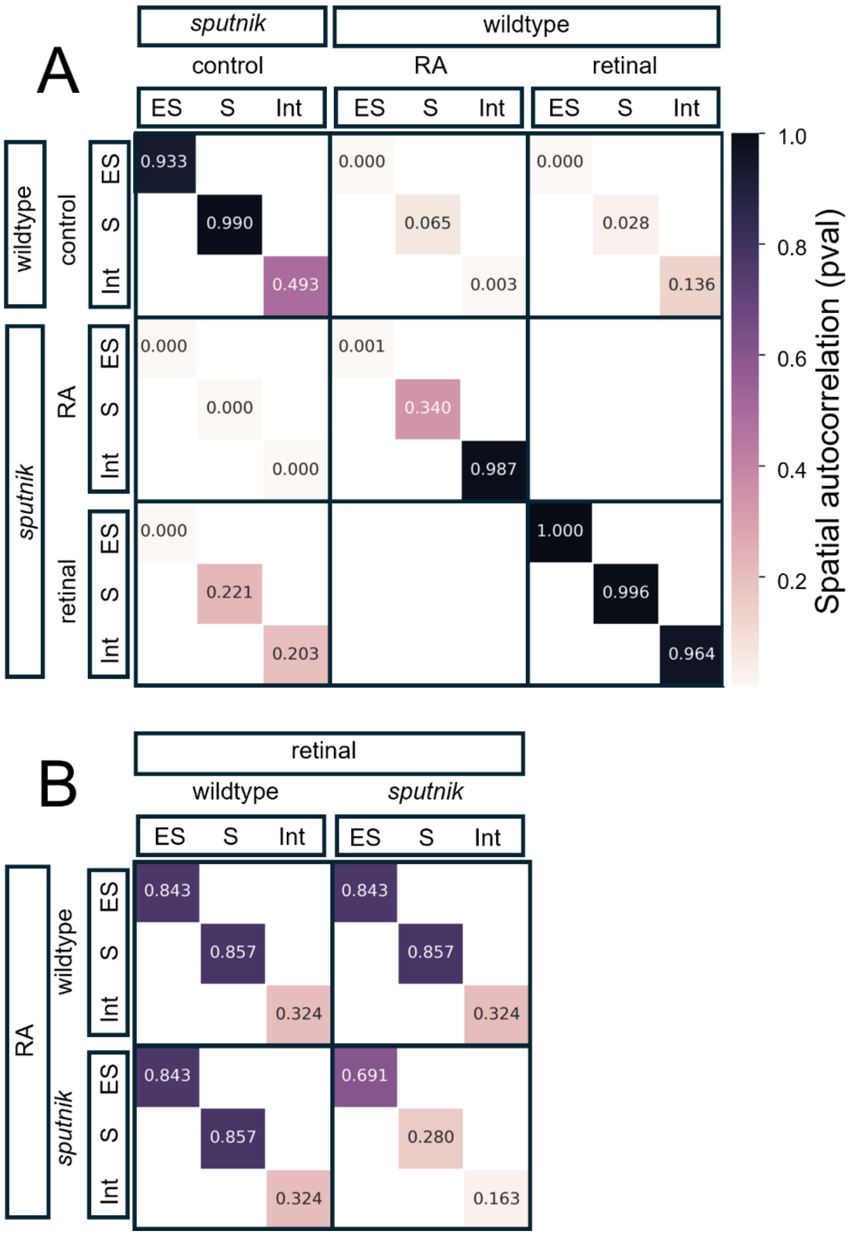


**Figure S6. Spatial autocorrelation analysis confirms differences in zonal patterning across conditions.**

Heatmap of spatial autocorrelation between indicated pairs. To determine overlap, nearest neighbor analysis was performed (k=7). Probabilities (pval) were generated by bootstrap analysis.
